# Cancer-cell-derived cGAMP limits the activity of tumor-associated CD8^+^ T cells

**DOI:** 10.1101/2024.10.28.620471

**Authors:** Michael Herbst, Hakan Köksal, Silvan Brunn, Dominik Zanetti, Ioana Domocos, Viola De Stefani, Paulo Pereira, Marc Nater, Virginia Cecconi, Maries van den Broek

## Abstract

Using a mouse tumor model with inducible cancer-cell-intrinsic cGAS expression, we show that cancer-cell-derived cGAMP is essential and sufficient to trigger a sustained type I interferon response within the tumor microenvironment. This led to improved CD8^+^ T cell-dependent tumor restriction. However, cGAMP limits the proliferation, survival, and function of STING-expressing but not of STING-deficient CD8^+^ T cells. *In vivo*, STING deficiency in CD8^+^ T cells enhanced tumor restriction. Consequently, cancer-cell-derived cGAMP both drives and limits the anti-tumor potential of CD8^+^ T cells. Mechanistically, T cell-intrinsic STING is associated with pro-apoptotic and antiproliferative gene signatures. Our findings suggest that STING signaling acts as a checkpoint in CD8^+^ T cells that balances tumor immunity.

**Graphical abstract:** 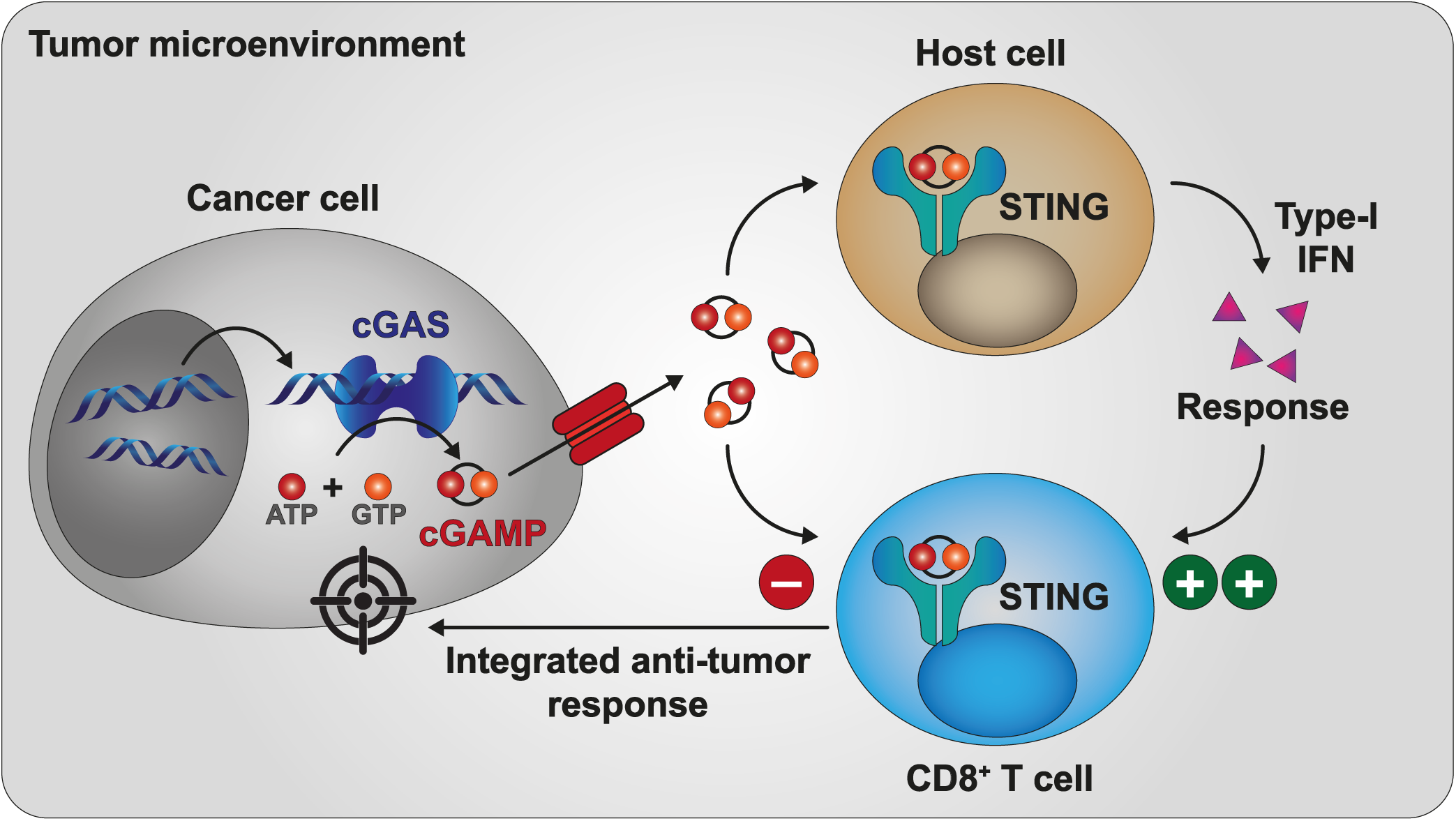

## INTRODUCTION

The presence of type I interferons (IFN-I) in the tumor microenvironment (TME) is essential for the generation of protective anti-tumor immune responses ^1–4^, largely through promoting antigen presentation and activating effector CD8^+^ T cells. IFN-I production occurs downstream of the cGAS-STING signaling pathway. This pathway is initiated when cytosolic double-stranded DNA (dsDNA) binds to cyclic GMP-AMP synthase (cGAS) ^5^, which catalyzes the formation of the second messenger cyclic GMP-AMP (cGAMP) ^6–10^. Subsequent binding of cGAMP to stimulator of interferon genes (STING) ^11^ results in the recruitment of TANK-binding kinase 1 (TBK1) ^12,13^ and phosphorylation of interferon regulatory factor 3 (IRF3) ^14,15^. Phosphorylated IRF3 then promotes transcription of IFN-I-encoding genes ^16^.

Whereas eukaryotic cells hardly contain cytosolic dsDNA in steady state, they contain substantial amounts under pathological conditions including viral infection, DNA damage, genomic instability, and cancer ^17^, which usually results in IFN-I production. However, cancer cells often lose their capacity to spontaneously produce IFN-I, suggesting that defects in the cGAS-STING pathway serve to escape immune surveillance ^18,19^. We and others showed that cGAMP can be transferred to other cells in the TME ^19–23^ to activate STING with subsequent IFN-I production ^19,24^. Despite the positive association between STING activation, IFN-I production and immune-mediated tumor control, STING signaling appears to play an ambiguous role in the TME. For example, intratumoral administration of a high amount of STING agonist resulted in the death of CD8^+^ T cells, and consequently compromised immune-mediated tumor control ^25^. STING signaling-induced T cell death was shown to rely on IRF3-driven upregulation of pro-apoptotic BH3 ^26^ and disrupted calcium homeostasis ^27^. Further, STING signaling in T cells reduces their proliferation ^28–30^, presumably in an NF-κB-dependent fashion ^29^. The functions of STING in murine T cells seem largely independent of IFN-I production ^31^. A study with primary human T cells showed that T cell-intrinsic STING activation enabled IFN-I production, but similarly impaired their function ^32^. Conversely, STING was required in adoptively transferred CD8^+^ T cells to maintain tumor-restricting efficacy ^33^. These contrasting findings suggest that deeper investigation into STING signaling in the TME and its effects on T cell responses is needed.

Most studies on STING signaling in T cells so far focused on CD4^+^ T cells and utilized artificial STING agonists ^26,28,29^, whereas only limited attention was given to CD8^+^ T cells in the TME. Whether tumor-derived cGAMP is sufficient to influence T cell-intrinsic STING signaling and how this affects the function of tumor-associated CD8^+^ T cells is unclear. The concept of enhancing adaptive immune responses by artificial STING agonists received considerable attention in the context of cancer ^34^. However, the detailed consequences of the transfer of cGAMP within the TME remain poorly understood.

Here, we show that cancer-cell-derived cGAMP results in sustained production of IFN-I and improved tumor control. At the same time, however, the activity of STING-expressing tumor-specific CD8^+^ T cells is curbed. Mechanistically, T cell-intrinsic STING is associated with pro-apoptotic and antiproliferative gene signatures, such that STING-deficient CD8^+^ T cells outcompete STING-expressing CD8^+^ T cells *in vivo*. Our findings suggest that STING signaling acts as a checkpoint in CD8^+^ T cells that regulates tumor immunity.

## RESULTS

### Cancer-cell-intrinsic cGAS expression restricts tumor progression

To study the immunological consequences of cancer-cell-derived cGAMP in the TME, we generated murine Lewis lung carcinoma (LLC) cells that express doxycycline (DOX)-inducible cGAS called LLC^cGAStetON^ (**Figure S1A**). Parental LLC cells do not express cGAS and hence, don’t produce cGAMP ^19^. The addition of DOX to cultured LLC^cGAStetON^ cells increased cGAS (*Mb21d1*)-transcripts (**Figure S1B**) and cGAS protein (**Figure S1C**). DOX administration to C57BL/6 mice with subcutaneous LLC^cGAStetON^ tumors induced the expression of cGAS (**Figures 1A** and **1B**), thus validating the experimental system. Further, we detected an increased concentration of the second messenger cGAMP in lysates of LLC^cGAStetON^ cells *in vitro* (**Figure S1D**) and in tumors after induced cGAS expression (**Figure 1C**). This indicates that sufficient dsDNA is present in LLC^cGAStetON^ cells to support cGAS-dependent cGAMP production. Cancer cells can export cGAMP via different mechanisms ^20,21,35^. We confirmed the export of cGAMP by LLC^cGAStetON^ cells by measuring cGAMP in the supernatant (**Figure S1E**). While induced cGAS expression did not affect the cell expansion *in vitro* (**Figure S1F**), we observed a significant reduction of subcutaneous (s.c.) tumor growth in immunocompetent C57BL/6 mice (**Figures 1D** and **1E**). DOX administration did not affect the growth of parental LLC tumors (**Figure 1F**), indicating that reduced tumor progression is due to cancer-cell-intrinsic cGAS, which is in line with previously published work ^19,24^. Thus, LLC^cGAStetON^ cells can be induced to express cGAS and produce cGAMP, which restricts tumor progression.

**Figure 1.**
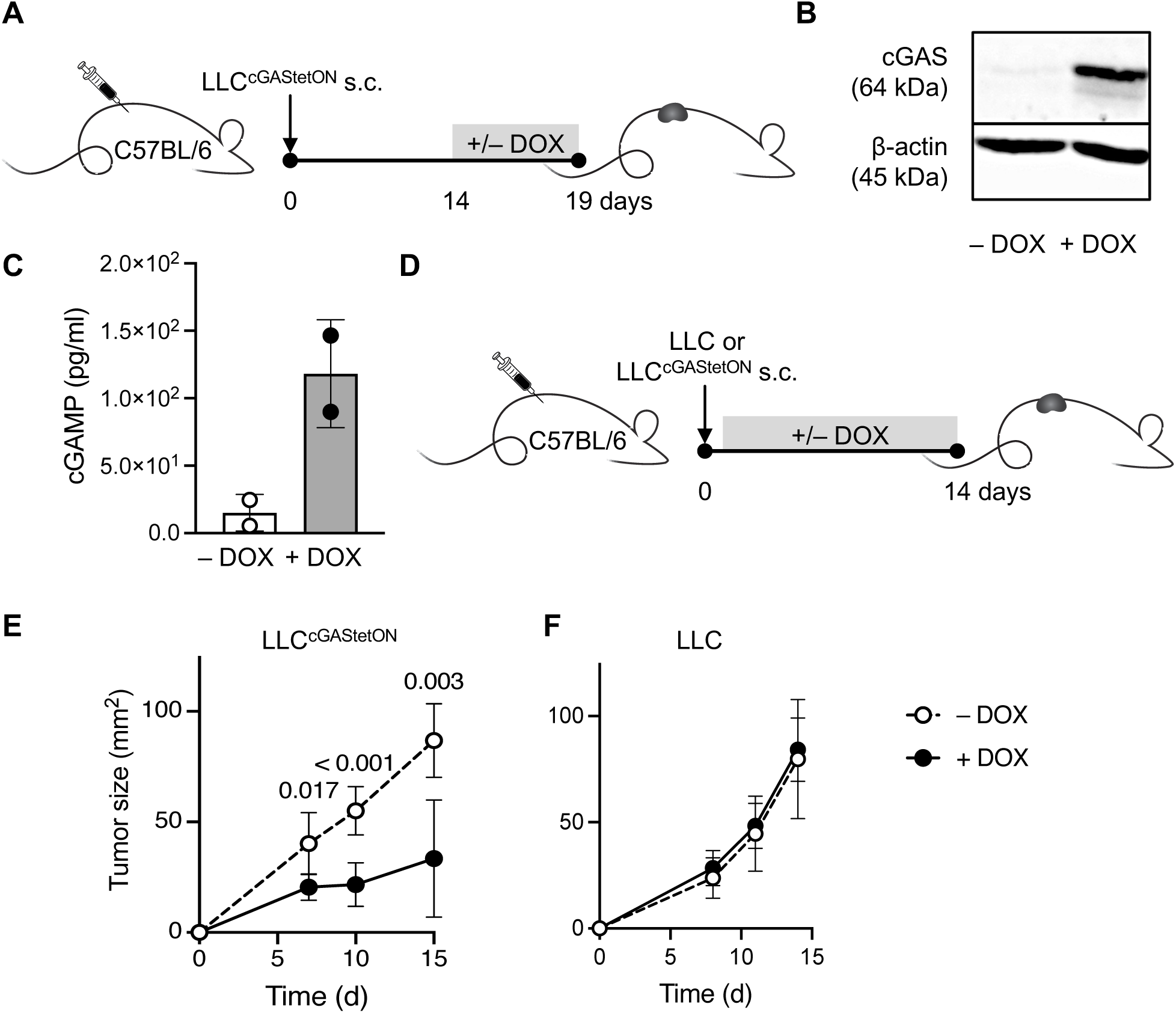
Induced expression of cancer-cell-intrinsic cGAS restricts tumor progression. **A** Experimental Setup. LLC or LLC^cGAStetON^ cancer cells (2 x 10^5^) were s.c. injected into C57BL/6 mice. Cancer-cell-intrinsic cGAS expression was induced 14 days later by providing 5 mg/ml DOX in sucrose-sweetened (5% w/v) drinking water. Control mice did not receive DOX. Tumors were isolated 5 days after induced cGAS expression. **B** Western blot detecting cGAS on *ex vivo* tumor samples. **C** 2′3′-cGAMP quantification (ELISA). Two technical replicates each. The same amount of total protein was assayed. **D** Experimental setup. LLC or LLC^cGAStetON^ cancer cells (2 x 10^5^) were s.c. injected into C57BL/6 mice. Cancer-cell-intrinsic cGAS expression was induced from the day of cancer cell injection as described in A. Control mice received no DOX. **E** Growth curve of LLC^cGAStetON^ tumors (n = 6). The graphs show mean ± SD. Each symbol represents an individual mouse. Statistical analysis: Unpaired two-tailed Student’s t-test per timepoint. **F** Growth curve of LLC tumors (n = 10). The graphs show mean ± SD. Each symbol represents an individual mouse. Statistical analysis: Unpaired two-tailed Student’s t-test per timepoint.

### Cancer-cell-derived cGAMP promotes a sustained IFN-I response and host STING-dependent tumor control

To investigate whether cancer-cell-derived cGAMP activates host STING, we quantified IFN-β, a member of the type I interferon (IFN-I) family, in LLC^cGAStetON^ tumors at different time points after induction of cGAS expression (**Figure 2A**). Whereas IFN-β was virtually absent before DOX administration, it was detectable after 16 hours of cGAS induction (**Figure 2B**). Using Mx1^gfp^ IFN-I response reporter mice, we found that both T and myeloid cells responded to IFN-I, with CD11c^+^ dendritic cells (DCs) and F4/80^+^ macrophages as major responders (**Figures 2C and S2A**). The IFN-I response remained consistent over time, suggesting that cGAMP-releasing tumors maintain an environment of sustained IFN-I response (**Figure 2D**). cGAMP and IFN-I are potent drivers of DC activation ^36^. We observed that the production of intratumoral cGAMP and IFN-I induced phenotypic changes in intratumoral DCs. Specifically, we saw an increase in the proportion of activated CD11b^+^Ly6C^+^ DCs (**Figures S2B-S2E**), a subset known to differentiate in tumors in response to tumor cell death ^37^.

**Figure 2.**
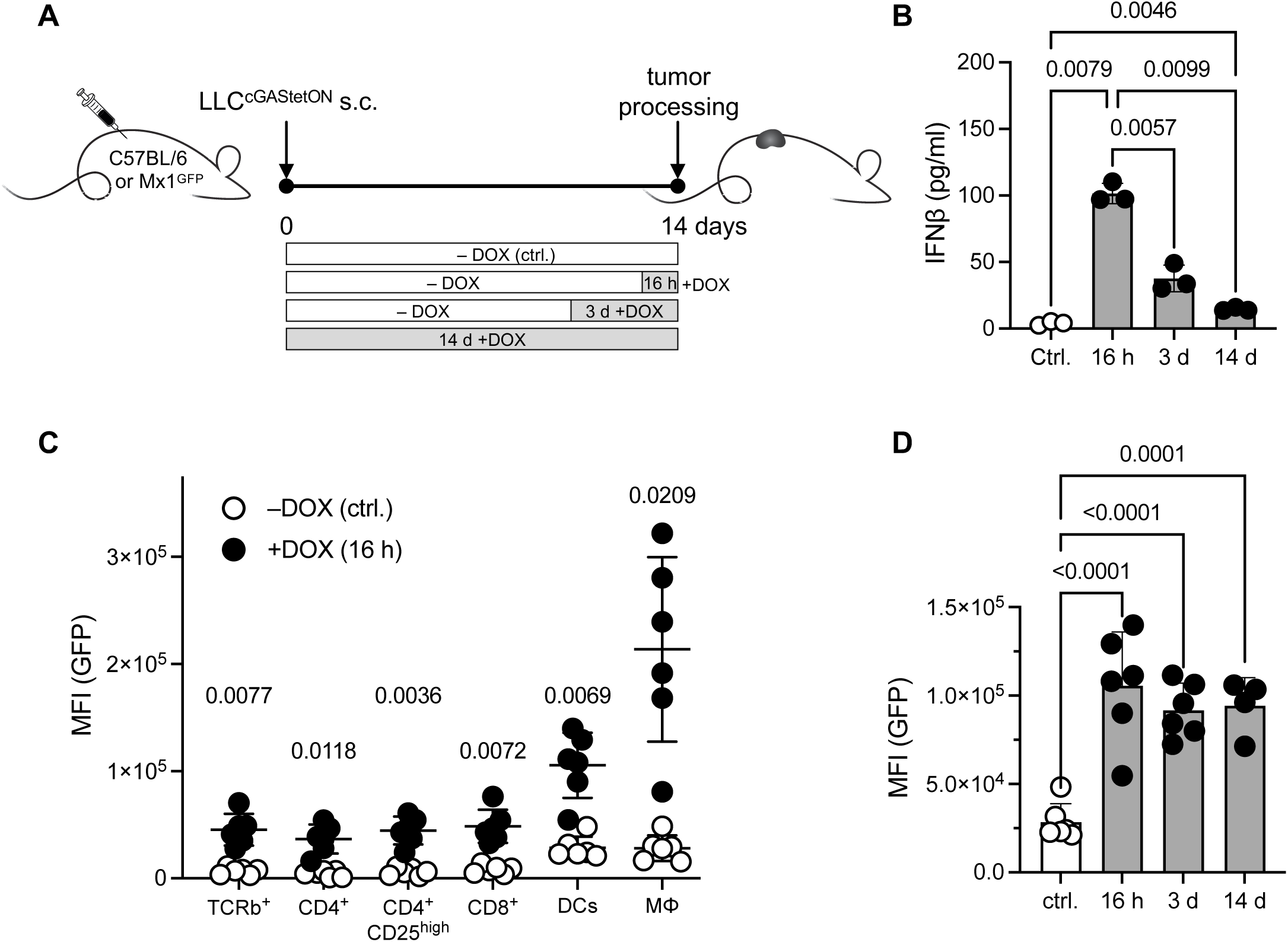
Cancer-cell-intrinsic cGAS promotes a sustained IFN-I response in the tumor microenvironment. **A** Experimental setup. LLC^cGAStetON^ cells (2 x 10^5^) were s.c. injected into C57BL/6 or Mx1^gfp^ IFN-I-response reporter mice. cGAS expression was induced by DOX during indicated periods. Control mice received no DOX. **B** Quantification of IFN-β in tumor lysates after periods of induced cGAS expression (n = 3 per group). Each symbol represents an individual mouse. Statistical analysis: One-way ANOVA with Welch’s correction. **C** IFN-I response in intratumoral cell populations measured in Mx1^gfp^ reporter mice 16 h after induction of cGAS expression (n = 6 per group). Each symbol represents an individual mouse. Statistical analysis: Two-way ANOVA. **D** Comparison of IFN-I-response in tumor derived DCs after induction of cGAS expression (n = 4-6 mice per group). Each symbol represents an individual mouse. Male and female mice were used. Statistical analysis: One-way ANOVA.

Since LLC cells express STING ^19^, we tested whether tumor cells contribute to IFN-I production via intrinsic STING engagement. An ELISA showed detectable IFN-β in LLC^cGAStetON^ cells + DOX *in vitro* (**Figure S2F**). Using CRISPR/Cas9, we generated LLC^cGAStetON-ΔSTING^ cells (**Figures S2G and S2H**). While LLC^cGAStetON-ΔSTING^ cells produced less IFN-β *in vitro*, the response to IFN-I *in vivo* was virtually unchanged (**Figures S2I and S2J**), suggesting that cancer-cell intrinsic STING signaling does not contribute to IFN-I in the tumor.

We introduced ovalbumin (OVA) in LLC^cGAStetON^ cells to allow monitoring of tumor-specific CD8^+^ T cells in future experiments and confirmed that induction of cGAS expression in LLC^cGAStetON-OVA^ tumors reduced tumor progression (**Figures S2K and S2L**). We verified that the improved control of cGAS-expressing tumors depends on host STING by using STING-deficient hosts (**Figures S2K and S2L**). STING deficiency in cancer cells did not impair tumor control (**Figures S2M and S2N**), providing further evidence for the importance of host STING. Collectively, our data suggest that cancer-cell-derived cGAMP promotes a sustained and host STING-dependent IFN-I response in the TME.

### STING-dependent tumor control depends on CD8^+^ T cells

Because IFN-I enhances antigen (cross-)presentation and the recruitment of CD8^+^ T cells to the tumor ^2,3,38^, we analyzed their infiltration in established tumors (**Figure S3A**). Already 5 days after induction of cGAS expression, we observed significantly more CD8^+^ T cells per gram of tumor (**Figure S3B**). To assess whether CD8^+^ T cells contribute to reduced tumor growth after induction of cancer-cell-intrinsic cGAS, we depleted CD4^+^ or CD8^+^ T cells starting at the time of DOX administration until the endpoint (**Figure 3A**). We found that the depletion of CD8^+^ but not CD4^+^ T cells abrogated the DOX-induced reduction of tumor growth (**Figures 3B** and **3C**). Depletion of CD4^+^ or CD8^+^ T cells was confirmed by flow cytometry (**Figure S3C**).

**Figure 3.**
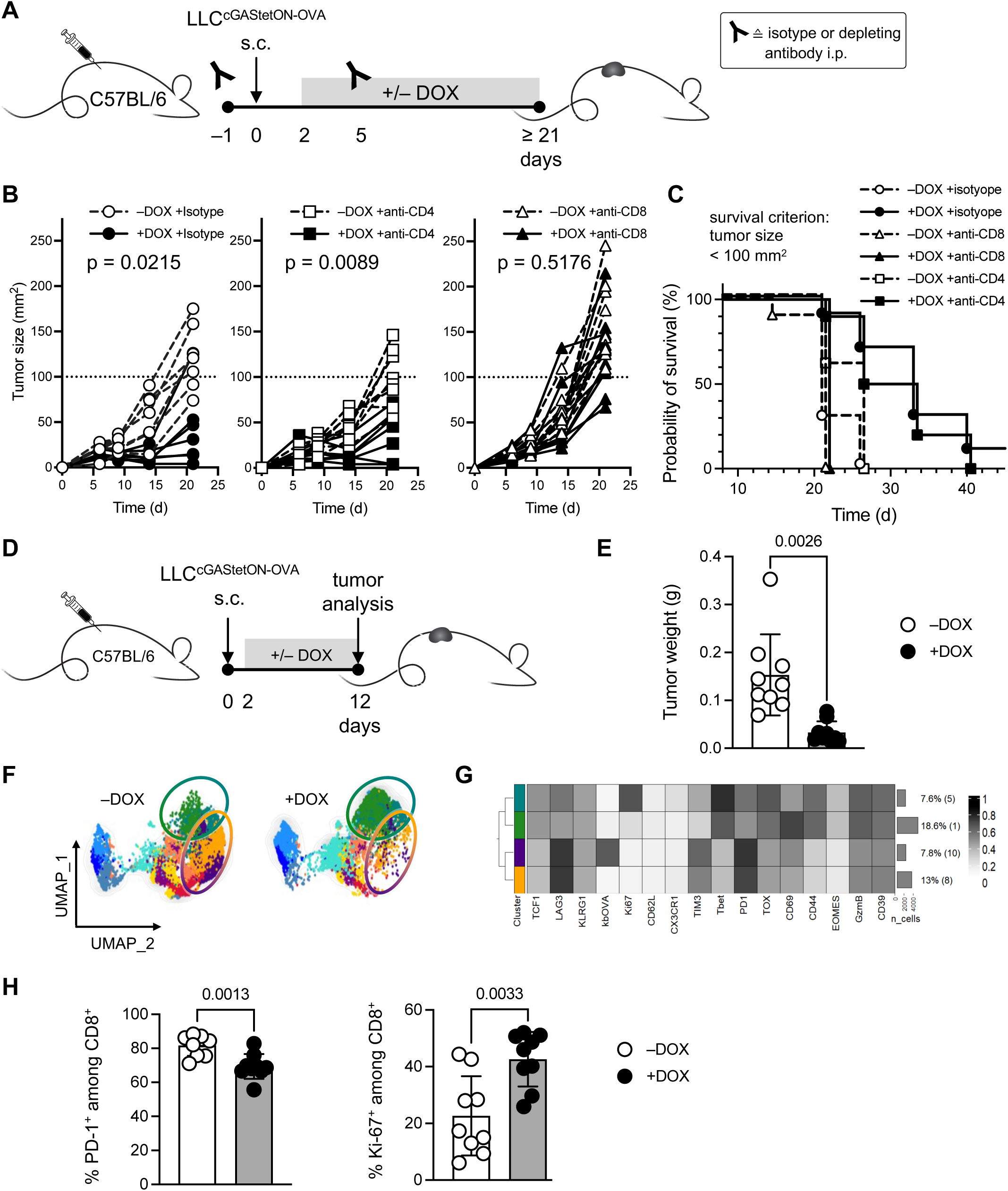
Induced expression of cancer-cell-intrinsic cGAS restricts tumor progression in a CD8^+^ T cell-dependent fashion. **A** Experimental setup for B and C. LLC^cGAStetON-OVA^ cells were s.c. injected into C57BL/6 mice. cGAS expression was induced by DOX from day 2 onwards until the endpoint. Control mice received no DOX. On day –1 and +5 relative to injection of cancer cells, mice received CD4- or CD8-depleting antibodies, or isotype as control. Each group consisted of 7-10 mice per group. **B** Tumor growth curves. Each symbol represents an individual mouse. Statistical analysis: Two-way ANOVA for data from day 21. **C** Survival curves. Death event is defined as tumor size ≥ 100 mm^2^. **D** Experimental setup for E-H. LLC^cGAStetON-OVA^ cells were s.c. injected into C57BL/6 mice. Cancer-cell-intrinsic cGAS expression was induced by DOX from day 2 onwards until the endpoint. Control mice received no DOX. **E** Tumor weights at the endpoint (day 12). Each symbol represents an individual mouse. Statistical analysis: Two-tailed t-test with Welch’s correction. **F** UMAP dimensionality reduction representation overlaid with identified clusters per condition (+/– DOX). **G** Heatmap of relative marker expression per identified T cell cluster. Partial figure shown, refer to supplement for full figure. **H** Frequency of PD-1^+^ and Ki-67^+^ cells among CD8^+^ T cells per condition. Each symbol represents an individual mouse. Statistical analysis: Two-tailed t-test with Welch’s correction.

Next, we investigated the influence of cGAMP and IFN-I on tumor-associated CD8^+^ T cells. Using the LLC^cGAStetON-OVA^ model we first confirmed that induced cGAS expression reduced tumor progression (**Figures 3D, 3E and S3D**). Clustering analysis (Rphenograph) of flow cytometry data collected 10 days after DOX administration identified 12 CD8^+^ T cell clusters. Identified clusters were projected on UMAP (Uniform Manifold Approximation and Projection) plots (**Figures 3F and S3E-S3G**). We found a higher frequency of terminally differentiated T cells (PD-1^high^, TIM3^high^, LAG-3^high^) in –DOX tumors and more active effector T cells (PD-1^+^, TIM3^low^, LAG-3^low^, GzmB^high^) in the +DOX condition (**Figures 3F** and **3G**). The overall proportion of PD-1^+^ CD8^+^ T cells was higher in the –DOX tumors, while the proportion of proliferating Ki-67^+^ CD8^+^ T cells was higher in the +DOX condition (**Figure 3H**). Together, these data suggest that a cGAMP- and IFN-I-rich TME supports the proliferation of infiltrated T cells while restricting their terminal differentiation.

Genotoxic treatments increase the concentration of cytoplasmic DNA ^39–43^, increase the amount of IFN-I and help to promote CD8^+^ T cell-dependent tumor control ^1,19,44^. We therefore proposed that radiotherapy is more efficient in cGAMP-producing LLC^cGAStetON^ tumors. To test this hypothesis, we treated mice with LLC^cGAStetON^ tumors with a single dose of 20 Gy (**Figure S3H**) as described ^44,45^. While the known effects of radiotherapy – reduction of tumor size and an increased number of CD8^+^ in the tumor – were detected, these effects were significantly more pronounced in cGAS-expressing LLC^cGAStetON^ tumors (**Figures S3I and S3J**). This suggests that cGAS activation and downstream production of cGAMP and IFN-I can be further amplified in our model.

### STING signaling in CD8^+^ T cells limits their activity *in vitro*

Production of cGAMP in the TME supports immune-mediated tumor restriction, which requires STING activation in host cells (**Figures S2K and S2L**) ^19,24^. cGAMP can be transferred to cells in the TME, and both cell type-specific and non-specific mechanisms have been suggested ^19,20,22,23,35,46–49^. We hypothesize that cancer-derived cGAMP directly influences CD8^+^ T cell behavior in tumors through STING activation. While *in vitro* studies showed that STING activation reduces proliferation and increases cell death of mostly CD4^+^ T cells ^26,28,29^, the overall consequences for CD8^+^ T cells are less clear.

To assess the functional effects of direct STING activation in CD8^+^ T cells, we established an *in vitro* assay in which TCR- and STING-signaling are concomitantly activated in OT-I T cells using the natural STING ligand cGAMP. Specifically, we cultured purified STING-proficient (STING^+/+^) or -deficient (STING^-/-^) OT-I CD8^+^ T cells with their cognate antigen (SIINFEKL) plus STING^-/-^ splenocytes as antigen-presenting cells. To induce STING signaling in OT-I cells, we added cGAMP to the cultures (**Figure 4A**). Addition of cGAMP significantly reduced multiple features associated with CD8^+^ T cell activation, including proliferation (Ki-67) and the expression of granzyme B (GzmB) in STING^+/+^ but not in STING^-/-^ OT-I CD8^+^ T cells (**Figures 4B-4E**). Further, exogenously added cGAMP resulting in STING signaling in CD8^+^ T cells reduced their viability (**Figure 4F**).

**Figure 4.**
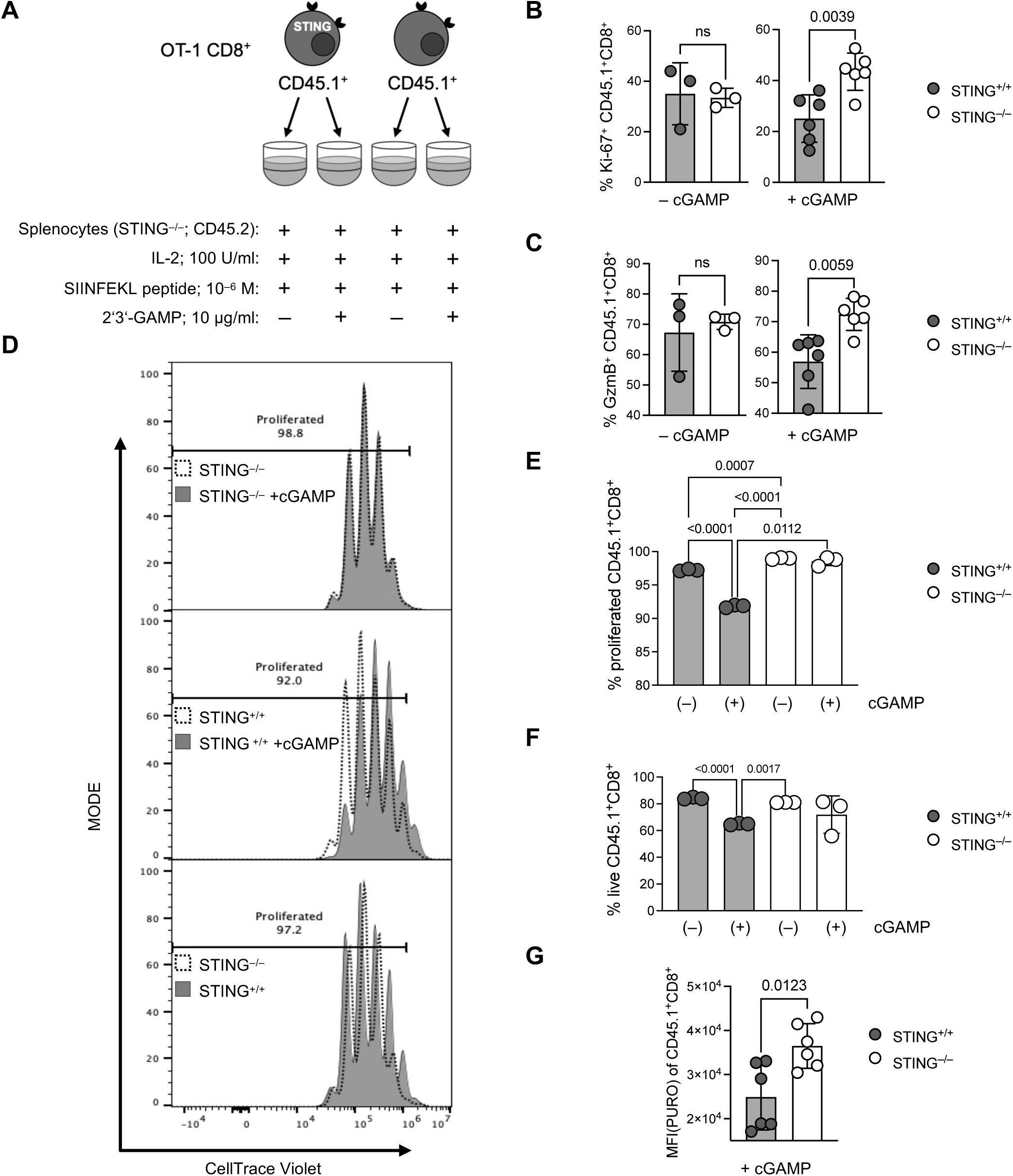
Concomitant TCR and STING signaling limits proliferation and metabolic activity of CD8^+^ T cells *in vitro*. **A** Experimental setup. Either 5 x 10^4^ CD45.1^+^ STING-proficient (STING^+/+^) or 5 x 10^4^ CD45.1^+^ STING-deficient (STING^-/-^) OT-I CD8^+^ T cells were stimulated with 10^-6^ M SIINFEKL and 2 x 10^5^ CD45.2^+^ splenocytes as antigen-presenting cells in the presence or absence of 10 µg/mL added cGAMP. For D-F, OT-I T cells were labeled with CellTrace Violet before culture start. After 48 h (B, C, G) or 60 h (D-F), cells were collected and analyzed by flow cytometry after gating on single, live CD45.1^+^ CD11b^-^ CD8^+^ cells. **B** Frequency of Ki-67^+^ CD45.1^+^ CD8^+^ of total cells. **C** Frequency of GzmB^+^ CD45.1^+^ CD8^+^ of total cells. **D** Representative proliferation curves of CellTrace violet-labelled OT-I cells. **E** Percentage of cells that underwent at least one proliferation cycle. **F** Frequency of live (LiveDead-NIR negative) CD8^+^ T cells. Cells gated on single CD45.1^+^ CD11b^-^ CD8^+^. **G** Incorporated puromycin (PURO) as measure for metabolic activity in CD45.1^+^ CD8^+^ cells. Experimental groups in B, C and G were statistically compared using two-tailed t-test with Welch’s correction. Experimental groups in E and F were statistically compared using one-way ANOVA. Data from 3 biological replicates (spleens) are shown (B, C, E-G). ns, not significant.

Because there is evidence for a connection between STING activation and metabolic fitness of T cells ^30,32^, we investigated the effect of exogenous cGAMP and consecutive STING signaling on the overall metabolic activity of cultured OT-I cells using the incorporation of puromycin (PURO) as a proxy ^50^. Again, exogenously added cGAMP reduced the metabolic activity of STING^+/+^ but not of STING^-/-^ CD8^+^ T cells (**Figure 4G**). Taken together, our results indicate that STING signaling in CD8^+^ T cells diminishes their proliferation, metabolic activity, and survival *in vitro*.

### STING signaling in CD8^+^ T cells limits their activity *in vivo*

After showing that STING signaling in CD8^+^ T cells reduces their activity *in vitro*, we next performed a series of *in vivo* experiments. First, we investigated the growth of cGAS-expressing tumors using CD4^Cre^STING^flox^ mice that lack STING specifically in T cells ^33,51^. Conditional STING deficiency was confirmed by flow cytometry (**Figure S4A**). T cell percentages in the thymus and spleen of CD4^Cre+/–^STING^flox^ and CD4^Cre-/-^STING^flox^ littermates were comparable ^33^ (**Figure S4B**). We induced cGAS expression in LLC^cGAStetON-OVA^ tumor-bearing mice (**Figure 5A**) and found that mice with T cell-specific STING deficiency had significantly smaller tumors than littermate controls (**Figures 5B** and **5C**). We conclude that STING in CD8^+^ T cells limits their efficacy in restricting tumors.

**Figure 5.**
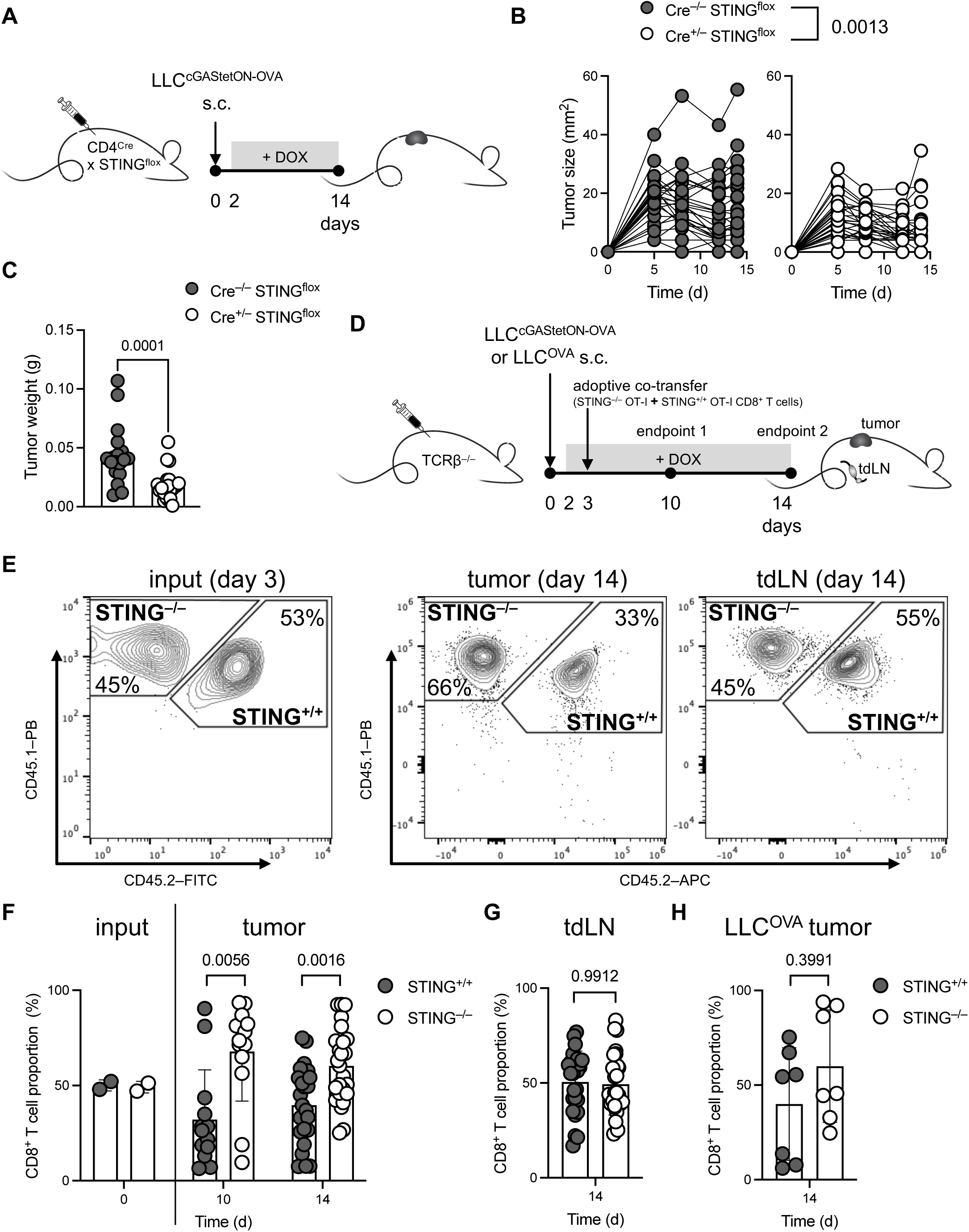
STING-deficient CD8^+^ T cells outcompete STING-proficient CD8^+^ T cells in the tumor. **A** Experimental setup for B and C. LLC^cGAStetON-OVA^ cells were s.c. injected into CD4^Cre+/–^STING^flox^ or CD4^Cre-/-^STING^flox^ littermates. cGAS expression was induced by DOX from day 2 onwards until the endpoint. **B** Tumor growth curves. Each symbol represents an individual mouse. Data from two pooled independent experiments with a total of 26-29 male and female mice. Statistical analysis: Two-way ANOVA. **C** Tumor weight at the endpoint. Statistical analysis: Two-tailed t-test. **D** Experimental setup for E-H. LLC^OVA^ or LLC^cGAStetON-OVA^ cells (3 x 10^5^) were injected s.c. into male and female TCRβ^-/-^ mice. cGAS expression was induced by DOX from day 2 onwards until the endpoint. On day 3, 1 x 10^3^ CD45.1/45.1 STING^-/-^ OT-I plus 1 x 10^3^ CD45.1/45.2 STING^+/+^ OT-I T cells were co-injected via the tail vein. The proportion OT-I cells was quantified in tumor and tumor-draining lymph nodes (dLNs) by flow cytometry at the endpoints. **E** Proportion of STING^-/-^ and STING^+/+^ OT-I cells at time of injection (day 0, left panel). Cells gated on single, live CD8^+^ cells. Representative examples for the frequency of STING^-/-^ and STING^+/+^ OT-I cells in tumor (middle panel) and tdLN (right panel) 11 days after adoptive transfer. Cells were gated on single, live CD45^+^ CD11b^-^ TCRβ^+^ CD8^+^ cells. **F** Proportion of STING^-/-^ and STING^+/+^ OT-I T cells at time of injection (day 0) and in LLC^cGAStetON-OVA^ tumors 7 and 11 days after transfer. **G** Proportion of STING^-/-^ and STING^+/+^ OT-I T cells in tdLNs 11 days after transfer. **H** Proportion of STING^-/-^ and STING^+/+^ OT-I T cells in LLC^OVA^ control tumors (no cGAS expression) 11 days after transfer. Experimental groups (F-H) were statistically compared using two-way ANOVA. Each symbol represents an individual mouse. Data from two pooled independent experiments with a total of 7-26 male and female mice. Data are expressed as mean ± SD.

Next, we directly compared the behavior of tumor-reactive STING^-/-^ and STING^+/+^ CD8^+^ T cells within an identical TME. Based on the STING-dependent reduction of proliferation and survival of CD8^+^ T cells described above (**Figure 4**), we hypothesized that STING^-/-^ T cells would outcompete STING^+/+^ T cells in a cGAMP-rich TME. To test this, we adoptively co-transferred naïve STING^-/-^ (CD45.1/1) and STING^+/+^ (CD45.1/2) OT-I CD8^+^ T cells into tumor-bearing, TCRβ^-/-^ (CD45.2/2) hosts that lack endogenous T cells but are otherwise immunocompetent (**Figure 5D**). We transferred equal numbers of STING^-/-^ or STING^+/+^ CD8^+^ T cells (**Figures 5E** and **5F**). After 10-14 days, we detected significantly more STING^-/-^ than STING^+/+^ OT-I cells in the tumors, whereas we found similar numbers of STING^-/-^ and STING^+/+^ OT-I cells in the tumor-draining lymph nodes (tdLN) (**Figures 5E-5G and S4C**). These differences could be explained by a local action of cGAMP, that seemed to be restricted to the tumor site. As a control, we included mice with LLC^OVA^ tumors that cannot produce cGAMP and found no differences in T cell frequency in tumor and tdLNs (**Figures 5H and S4D**). Taken together, our results suggest that cancer-cell-derived cGAMP restricts CD8^+^ T cells via T cell-intrinsic STING activation in the tumor.

### STING activity in CD8^+^ T cells promotes their apoptosis and reduces their proliferation in a cGAMP-rich environment

To understand the differences between STING-deficient and -proficient intratumoral CD8^+^ T cells, we again co-transferred STING^-/-^ (CD45.1/1) and STING^+/+^ (CD45.1/2) CD8^+^ T cells into TCRβ^-/-^ (CD45.2/2) hosts bearing cGAS-expressing LLC^cGAStetON-OVA^ tumors. Six days later, we sorted the transferred CD8^+^ T cells from the tumors and performed single-cell RNA sequencing (**Figures 6A and S5A**). Co-transfer and reisolation allowed the comparison of STING-deficient and -proficient CD8^+^ T cells from the same TME. As expected, T cells isolated from tumors with a cGAMP-rich TME (+DOX) showed a pronounced interferon response signature compared to – DOX (**Figures S5B and S5C**), thus validating STING activation in the TME. Clustering based on UCell ^52^ scoring indicated fewer exhausted CD8^+^ T cells in cGAS-expressing tumors (**Figure S5D**), which goes in line with our previous observations (**Figures 3F** and **3G**). Comparing STING^-/-^ and STING^+/+^ CD8^+^ T cells from the same TME showed overall similar results regarding the frequency of 11 identified cell clusters (**Figure 6B**). A gene set enrichment analysis (GSEA), however, revealed significant changes in different pathways. Genes associated with apoptosis were downregulated in STING^-/-^ T cells, suggesting reduced apoptotic processes in the absence of STING (**Figure 6C**). This observation was supported by an overall higher differential expression of Caspase 3 (*Casp3*), a key gene in the apoptosis pathway of T cells ^53^. The association between STING expression and apoptosis aligns with our *in vitro* data where we found reduced cell survival of STING^+/+^ CD8^+^ T cells when incubated with cGAMP (**Figure 4F**) and with previous studies that link STING to apoptosis in T cells ^26,28^. Further, the GSEA pointed towards reduced inflammatory responses and signaling via NFκB in STING^-/-^ CD8^+^ T cells. The fact that NF-κB is a key downstream target of STING may explain this finding. Because NF-κB-dependent antiproliferative effects have been suggested for T cells ^29^, we compared the proliferating cell fraction (clusters 3 and 7) of STING^+/+^ and STING^-/-^ CD8^+^ T cells (**Figure S5D**). STING^-/-^ T cells expressed significantly higher *Mki67* compared to STING^+/+^ T cells, indicating reduced proliferation when STING is present. In addition, *Ifi208* (an IFN-response gene), and *Camta1* and *Grm7* (involved in calcium homeostasis ^54,55^) were highly expressed in STING^-/-^ T cells (**Figure S5E**). A previous study indicated that STING engagement influences calcium signaling in T cells and leads to apoptosis ^27^. Of note, we did not detect intrinsic IFN-β gene expression in reisolated CD8^+^ T cells (**Figure S5F**).

**Figure 6.**
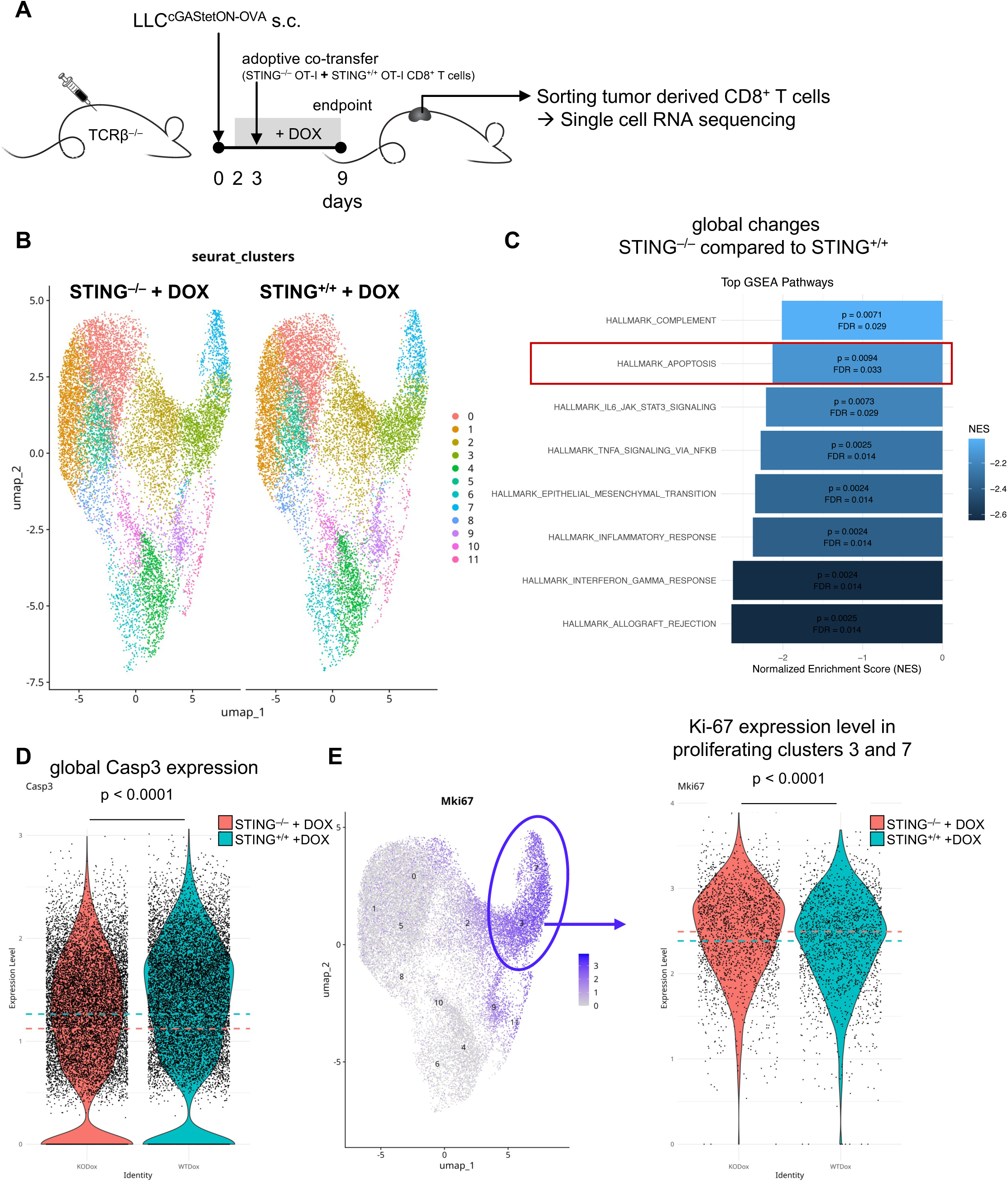
STING expression in CD8^+^ T cells promotes their apoptosis and reduces their proliferation in a cGAMP-rich environment. **A** Experimental setup. LLC^cGAStetON-OVA^ cells (3 x 10^5^) were injected TCRβ^-/-^ mice. cGAS expression was induced by DOX from day 2 onwards until the endpoint. On day 3, 5 x 10^3^ CD45.1/45.1 STING^-/-^ OT-I plus 1 x 10^3^ CD45.1/45.2 STING^+/+^ OT-I T cells were co-injected via the tail vein. On day 9, CD8^+^ T cells were sorted from tumors and subjected to single-cell whole transcriptome analysis. **B** UMAP representation of 12 (0-11) identified Seurat clusters. **C** Gene set enrichment analysis (GSEA) using Molecular Signatures Database (MSigDB) gene sets. **D** Global expression comparison of Casp3. **E** Ki67 expression in subsetted proliferating clusters 3 and 7, shown in the feature plot (left panel). Data are shown from 6 individually hashed mouse/tumor samples. Experimental groups were statistically compared using two-way ANOVA followed by Tukey’s Honest Significant Difference (HSD) test.

In conclusion, when CD8^+^ T cells are exposed to a cGAMP-rich environment, such as within the TME, STING activation promotes their apoptosis, triggers an inflammatory response, and suppresses their proliferation. Consequently, STING-deficient CD8^+^ T cells mediate superior protective anti-tumor immunity. Our findings suggest that STING expression in CD8^+^ T cells might act as an additional immune checkpoint in tumors.

## DISCUSSION

STING-signaling is as a critical pathway for the regulation of immune responses across various cell types, including stromal ^12^, endothelial ^56,57^, epithelial ^58^, myeloid ^2,36,59^, and lymphoid cells ^29,60,61^. Consistent with previous observations ^19,24^, we showed that host STING is essential for tumor restriction in a model with inducible cancer-cell-intrinsic cGAS expression. STING activation was driven by cGAMP-transfer from cancer cells to other cells in the tumor microenvironment and led to CD8^+^ T cell-mediated tumor restriction. We demonstrate that cancer-cell-derived cGAMP exerts both beneficial and inhibitory effects on T cell mediated tumor control. There is ample evidence that the activation of STING by cGAMP or artificial STING agonists induces protective anticancer immunity responses, restricts tumor progression, and synergizes with immune checkpoint inhibition ^36,57,59,62–64^. However, clinical phase I/II trials combining immunotherapy with STING agonists have shown varying efficacy ^34^, suggesting that a more precise understanding of the mechanisms underlying STING activation in tumors is needed.

Beyond inducing IFN-I, STING activation exerts IFN-I-independent effects, such as promoting different forms of cell death ^65^. We found that *in vitro* activation of CD8^+^ T cells by their cognate antigen was inhibited by addition of exogenous cGAMP. This finding aligns with previous studies showing that concurrent TCR and STING activation impairs T cell proliferation ^28,29^. Similarly, metabolic impairment in T cells after TCR and STING activation, presumably due to impaired mTORC1 signaling in an IRF3/IRF7-dependent manner, has been reported ^30,32^. We observed STING-dependent inhibition of CD8^+^ T cells also in the tumor microenvironment *in vivo*. Our finding that STING-deficient CD8^+^ T cells outcompeted STING-proficient ones in the same tumor shows that STING activation inhibits CD8^+^ T cells in a cell-intrinsic manner, independent of other cells. Single-cell RNA sequencing data indicated an increased apoptosis-related gene-signature and decreased proliferation-related gene expression in CD8^+^ T cells when STING was present. We did not detect *Ifnb* transcripts in CD8^+^ T cells, supporting earlier findings that murine T cells can function independently of IFN-I production ^31^ and thus seem to behave differently from human T cells in this aspect ^32^.

While some studies also reported negative effects of STING activation on T cell survival ^26,28,29^, other literature presents conflicting results. For example, Li and colleagues suggested that CD8^+^ T cell-intrinsic STING supports tumor-specific functions ^33^. Along the same line, STING activation in CD4^+^ T cells enhanced antitumor responses through supporting differentiation into effector cells ^66^. A recent study using conditional STING deficiency in T cells suggested slower tumor progression, which was explained by STING activation-dependent expansion of CD4^+^FoxP3^+^ regulatory T cells ^51^. Possible explanations for the above-mentioned discrepancies may be differences in experimental models and readouts. For instance, most studies have used synthetic STING ligands like DMXAA ^67^, which may act differently compared to cGAMP, as it can exert additional functions beyond STING activation ^68^. Moreover, locally produced cGAMP may have different effects compared to systemic or intratumorally administered STING agonists.

We did not investigate cGAMP transfer mechanisms in the tumor microenvironment or an eventual role of cGAMP-hydrolyzing enzymes like ENPP1 ^46^. The strong IFN-I production in tumors following induced cGAS expression makes it unlikely that cGAMP-hydrolyzing enzymes play a significant role in our model. LRRC8c ^69^ and SLC19A1 ^22,23^ have recently been identified as cGAMP importers that can facilitate the uptake of cGAMP. It may be interesting to investigate the contributions of these transporters in modulating immune responses in different cell types within the tumor microenvironment, as well as the effect on immune-mediated tumor restriction.

Taken together, we have shown that cancer-cell-derived cGAMP modulates T cell activity, inducing both positive and inhibitory effects. Therapeutic strategies targeting STING should carefully balance innate immune activation with the protection of T cells from STING-mediated inhibition to maximize treatment efficacy. The inhibitory action of STING in CD8^+^ T cells could be a contributing factor to CAR-T cell dysfunction in chronic antigen exposure. Thus, it may be worth investigating whether STING-deficient CAR-T cells are more efficacious than STING-expressing counterparts. The fact that T cells retain STING expression throughout differentiation suggests a physiological role for STING as an immune checkpoint, possibly evolved to limit T cell overactivation in chronic inflammatory environments like cancer or persistent viral infection.

## Acknowledgments

This work was supported by the University Research Priority Program “Translational Cancer Research” (University of Zurich; MvdB), SKINTEGRITY.ch (University of Zurich; MvdB), the Hartmann-Müller-Foundation (MH), the Swiss National Science Foundation (310030_208145 MvdB), the Stiftung für Krebsbekämpfung Zurich (MvdB) and the Novartis Foundation for medical-biological Research (24A054; MvdB). The authors thank the personnel of the Laboratory Animal Services Center (LASC, University of Zurich) for expert animal care. We thank Tatiane Gorski (Cytometry Facility, University of Zurich) and Hubert Rehrauer (Functional Genomic Center Zurich) for help with scRNA-sequencing, and Ulrich Kalinke (Twincore, Hannover, Germany) and Hans Christian Probst (University of Mainz, Germany) for discussion.

## Author contributions

MH and MvdB conceived the experiments and wrote the manuscript; MH, HK, SB, DZ, ID and VDS performed the experiments. PP and VC performed sorting. MH, HK, and MN analyzed the data. MvdB secured funding; All the authors reviewed the results and approved the final manuscript.

## Ethical approval statement

Mouse experiments were performed according to Swiss cantonal and federal regulations on animal protection and approved by the cantonal veterinary office of Zürich under license numbers 37/2021 (33442) and 38/2021 (33443).

## STAR METHODS

### RESOURCE AVAILABILITY

#### Lead contact

Further information and requests for resources and reagents should be directed to and will be fulfilled by the lead contact, Maries van den Broek.

#### Materials availability

The materials generated for this study can be provided upon reasonable request.

#### Data and code availability

- All data reported in this paper will be shared by the lead contact upon reasonable request.
- This paper does not report an original code.
- Any additional information required to reanalyze the data reported in this paper is available from the lead contact upon reasonable request.

## EXPERIMENTAL MODEL DETAILS

### Mice

C57BL/6NRj mice (8-10 weeks old) were purchased from Janvier Labs. STING-deficient (STING^-/-^, C57BL/6J-*Sting1^gt^*/J) ^70^, STING^flox^ (B6;SJL-*Sting1^tm1.1Camb^*/J) ^71^ and Mx1^gfp^ (B6.Cg-*Mx1^tm1.1Agsa^*/J) ^72^ mice were purchased from Jackson Laboratory. *Tcrb*-deficient (B6.129P2-*Tcrb^tm1Mom^*/J) ^73^ and CD4^Cre^ (STOCK Tg(Cd4-cre)1Cwi/BfluJ) ^74^ mice were originally obtained from the Jackson Laboratory. *Tcrb*-deficient mice were provided by Annette Oxenius (ETH Zurich, Switzerland). CD4^Cre^ mice were provided by Burkhard Becher (University of Zurich, Switzerland). CD4^Cre^ x STING^flox^ and STING^-/-^ x OT-I ^75^ x Ly5.1 mice were bred in-house. All strains are on a C57BL/6 background. All mice were housed in individually ventilated cages with wood chip bedding and maintained in a temperature-controlled environment with a 12-hour light/dark cycle under specific pathogen-free (SPF) conditions at the facilities of the Laboratory Animal Services Center (LASC) at the University of Zurich. Mice had access to water (containing 0.5 ppm ClO_2_) and food *ad libitum*. All experiments were performed with 8-12 weeks-old female mice unless stated otherwise. Experiments were approved by the Cantonal Veterinary Office Zurich under the license numbers ZH37/2021 and ZH38/2021 and conducted according to federal and cantonal regulations.

### Cell lines and cell culture

LLC cells were purchased from ATCC (CRL-1642). MC38 cells were originally provided by Mark Smyth (QIMR Berghofer Medical Research Institute, Brisbane, Australia). Cell lines were tested negative by PCR for *Mycoplasma ssp.* in-house and for 18 additional mouse pathogens (IMPACT II Test, IDEXX Bioanalytics). Cancer cells were cultured in DMEM (Gibco) supplemented with 10% fetal bovine serum (Gibco), 100 U/mL penicillin, 100 μg/mL streptomycin (Sigma) and 2 mM L-glutamine (Gibco) at 37°C in a humidified atmosphere containing 5% CO_2_. For counting, cells were diluted 1:100 in 0.4 % Trypan Blue Solution (Gibco) and manually counted using a Neubauer-improved counting chamber. Where indicated, LLC^cGAStetON^ cells were cultured with 1 µg/mL DOX and counted after 1 to 4 days.

## METHOD DETAILS

### Modification of cell lines

For the generation of LLC^cGAStetON^ cells, murine cGAS-DNA was amplified from the vector pLenti-EF1α-Flag-mm-cGas using Fwd_ArvII-cGAS and Rev_cGAS-MluI primers (**Table S1**), while adding restriction sites. Amplified DNA was excised and purified from a 1% agarose gel using the NucleoSpin Gel & PCR Clean-up Mini kit (Macherey-Nagel). The target vector (pCW57-GFP-P2A-MCS) and cGAS amplicon were digested with AvrII and MluI-HF. Restricted fragments were purified from a 1% agarose gel and ligated with T4 DNA ligase (Thermo Fisher Scientific). Plasmids were amplified in heat shock-transformed (30 seconds at 42°C) T10 *Escherichia coli* that were grown on agar plates containing ampicillin for selection. Successful cloning was verified by Sanger sequencing of single colonies using the E.coli NightSeq service (Microsynth). Plasmid DNA was purified from liquid bacterium cultures using the ZymoPure II Plasmid Maxiprep kit (Zymo Research). Lentivirus for modification of LLC cells was generated with a second-generation lentiviral packaging system (pMD2.G and pCMV-dR8.91) in HEK 293T cells. A DNA:polyethylenimine (Polysciences) ratio of 1:3 was used for transfection. Lentiviral transduction was performed with 1:5 virus-dilution in DMEM supplemented with 8 µg/mL polybrene (Sigma-Aldrich), followed by 45 minutes of spinoculation at 37°C and 800 *g*. After 48 hours, 500 µg/mL neomycin (InvivoGen) was added to select transduced cells. The DOX-induced expression of cGAS was confirmed by Western blot.

For the generation of LLC^cGAStetON-OVA^ cells, virus generation and transduction were performed as described above using the pHR OVA/p2a/mCherry-CaaX (Addgene #113030) plasmid. Six days after transduction, cells were sorted for live (Zombie NIR^-^) mCherry^+^ cells.

For the generation of LLC^cGAStetON-ΔSTING^ cells, single guide (sg) RNAs targeting mouse STING (*Tmem173*) were designed using CRISPOR (http://crispor.gi.ucsc.edu)^76^ (**Table S1**). SgRNAs were cloned into pSpCas9(BB)-2A-GFP (PX458) (Addgene #48138) after BpiI-mediated restriction. Plasmid amplification and purification were performed as described above. Target cells were transformed using Lipofectamine 3000 (Invitrogen). After 2 days of culture, GFP^+^ cells were FACS sorted and expanded. Deficiency of STING was confirmed by Western blot and flow cytometry. The LLC^cGAStetON^ model provides significant advantages over stable knock-in/out models, as it avoids unintentional differences caused by the generation of distinct cell lines.

### *In vivo* experiments

Cultured cancer cells were harvested and suspended at 3 x 10^6^/mL in a 2:1 ratio of PBS:Matrigel (Corning). If not stated otherwise, 3 x 10^5^ cancer cells (100 µL) were injected subcutaneously (s.c.) into the right, previously shaved flank. Unless otherwise specified, cancer-cell-intrinsic cGAS expression was induced by an initial intraperitoneal (i.p.) injection of 50 µg/kg DOX in PBS and subsequent administration of 2 mg/mL DOX + 0.05 mg/mL sucralose in the drinking water for indicated periods. Groups without induction of cGAS expression received PBS i.p. and sucralose-containing (0.05 mg/mL) drinking water. The tumor size was measured in two dimensions (length and width) using a digital caliper.

For T cell depletion, 500 µg of CD4-depleting antibody (clone GK1.5; Rat IgG2a) or CD8-depleting antibody (clone YTS169.4; Rat IgG2a) was injected i.p. Anti-KLH antibody (clone LTF-2; BioXcell) served as an isotype control antibody. After 5 days a second injection was performed, and cell depletion was confirmed in the blood using a CyAn ADP (Beckman Coulter) flow cytometer. For survival studies, a death event was recorded when the tumor size reached 100 mm^2^.

For adoptive transfer, CD8^+^ T cells were purified from the spleen of age- and sex-matched OT-I mice using the EasySep Mouse CD8^+^ T Cell Isolation Kit (STEMCELL Technologies, Cat. #19853) according to the manufacturer’s instructions. The purity of enriched CD8^+^ T cells was assessed by flow cytometry and was > 95%. Purified CD8^+^ T cells were injected in 200 μL sterile PBS via the tail vein. The number of transferred cells is specified in the individual experiments.

For radiotherapy, tumor-bearing mice were randomized based on tumor size immediately before radiotherapy. Radiotherapy was performed as described ^45^. Briefly, mice were anesthetized by i.p. injection of 50 mg/kg ketamine and 10 mg/kg xylazine. Vitamin A eye ointment was applied to the eyes to prevent dryness during the procedure. Mice were secured in a lead cage to ensure localized irradiation of the tumor with a single dose of 20 Gy using an RS-2000 irradiation unit (Rad Source) with a dose rate of 1.81 Gy/min.

In experiments where cGAS expression was induced in established tumors (> day 5), mice were randomized into experimental groups with similar average tumor size and variance before the intervention. Tumors that were palpable but too small to reliably measure were assigned a size of 4 mm^2^.

### Tissue collection and processing

Tumors and tumor-draining lymph nodes (tdLNs) were excised and collected in RPMI containing 10% FCS. Tumors were manually cut into small pieces and subsequently digested in RPMI containing 10% FCS, 1 mg/mL collagenase IV (Thermo Fisher Scientific), and 50 µg/mL DNase I (Roche) for 45 minutes at 37°C on a rotating device. Digestion was stopped by the addition of 5 mL ice-cold PBS. Digested tumors were passed through a 70-µm cell strainer using the plunger of a 5-mL syringe. Cells were washed with PBS and pelleted by centrifugation at 350 *g* for 5 minutes. Red blood cells were lysed using RBC lysis solution (17 mM Tris pH 7.2, 144 mM NH_4_Cl) for 2 minutes at room temperature. tdLNs were directly passed through a 70-µm cell strainer and washed with complete RPMI. Blood was collected by submental bleeding in PBS containing 2 mM EDTA.

### *In vitro* experiments

Isolated CD8^+^ CD45.1^+^ OT-I were labeled with CellTrace Violet (Thermo Fisher Scientific) according to the manufacturer’s instructions. Fifty thousand labeled CD8^+^ CD45.1^+^ OT-1 STING-deficient or -proficient T cells were co-cultured with the cognate antigen (SIINFEKL), 2 x 10^5^ CD45.2^+^ STING-deficient splenocytes as antigen-presenting cells and 10 µg/mL cGAMP (or not). Cells were cultured in RPMI containing 10% FCS, 100 U/mL penicillin, 100 μg/mL streptomycin, and 100 U/mL IL-2 for the period indicated in the figures.

### Western blot

To extract proteins, cultured cells or tumor tissue were suspended in RIPA buffer (Thermo Fisher Scientific) containing proteinase inhibitor (cOmplete, Mini Protease Inhibitor Cocktail, Roche)). Cells were lysed by pipetting up and down, whereas a handheld homogenizer was used to lyse tumor tissue. Samples were centrifuged at 14,000 *g* for 15 minutes, and protein-containing supernatant was removed. The protein concentration was measured using the DC Protein Assay (Bio-Rad). A total of 20 µg protein was loaded on a 10% SDS PAGE gel. Electrophoresis was performed using a Bio-Rad system at 80 V during the stacking phase for 30 minutes, followed by 120 V for about 1 hour. Proteins were transferred onto a nitrocellulose membrane (GE Healthcare Life Science) with a current of 350 mA for 2 hours at 4°C. The membrane was blocked in 5% non-fat milk powder in Tris-buffered saline containing 0.05% Tween-20 (TBS-T) for 1 hour at room temperature or overnight at 4°C. Membranes were stained with primary antibodies against cGAS (D3O8O, Rabbit mAb #31659, Cell Signaling Technologies, 1:1,000), STING (D2P2F, Rabbit mAb #13647, Cell Signaling Technologies, 1:1,000 dilution) or β-actin (clone AC-15, mouse monoclonal #A5441, Sigma-Aldrich, 1:5,000 dilution) for 1 hour at room temperature. Primary antibodies were diluted in TBS-T containing 5% non-fat milk powder. Membranes were washed three times for 10 minutes in TBS-T and incubated with the secondary antibodies diluted 1:5,000 in TBS-T containing 5% non-fat milk powder for 1 hour at room temperature. HRP-goat-anti-mouse IgG (Dianova, #111-035-008) and HRP-goat-anti-rabbit IgG (Dianova, #111-035-045) were used as secondary antibodies. Membranes were again washed three times for 10 minutes in TBS-T, and a chemiluminescent reaction was started using the WesternBright ECL-HRP kit (Advansta, K-12045-D20). Protein bands were imaged with a Fusion Solo S imager (Vilber).

### Flow cytometry

Single-cell suspensions were incubated with anti-CD16/32 antibody in PBS for 5 minutes. Antibody staining of surface molecules was conducted in 50 µL antibody mix in PBS for 30 minutes at 4°C. For intracellular staining, cells were fixed and permeabilized using the Foxp3/Transcription Factor Staining Buffer Set (Invitrogen) for 45 minutes at 4°C according to the manufacturer’s instructions. The antibody mix for staining of intracellular molecules was prepared in 1X permeabilization buffer, and cell suspensions were stained with 50 µL overnight at 4°C. Samples were washed once with 1X permeabilization buffer and once with PBS before acquisition on a Cytek Aurora 5L spectral flow cytometer. Alternatively, LSRFortessa (BD Biosciences) or CyAn ADP (Beckman Coulter) devices were used.

Puromycin incorporation, a proxy for metabolic activity ^50^, was determined by flow cytometry. CD8^+^ T cells were resuspended in 100 µL RPMI containing 2% FBS and were rested at 37°C. After 15 minutes, 10 µL puromycin (10 µg/mL in DMSO) was added, and cells were incubated for an additional 40 minutes at 37°C. After incubation, the cells were washed with 150 µL ice-cold PBS. Subsequent staining for flow cytometry was performed as described above. Anti-puromycin antibody was added during the intracellular staining step at a dilution of 1:1,000.

FCS files were processed using FlowJo version 10.9. Data from gated populations of interest were exported as compensated FCS files. Preprocessed FCS files were imported into R version 4.2.2. Data were arcsinh transformed and raw marker intensities were quantile normalized to ensure consistent sample comparisons. Subsequent dimensionality reduction was performed using t-distributed Stochastic Neighbor Embedding (t-SNE) or Uniform Manifold Approximation and Projection (UMAP) ^77^. Cell clusters were identified using unsupervised Rphenograph-clustering^78^. Data were visualized using the R package ggplot2. Bioconductor version 3.16 was used for analysis.

### Single-cell RNA sequencing

Single-cell suspensions from tumors (*n* = 6 per condition) were labeled using the BD Single-Cell Multiplexing Kit (BD) and pooled according to their respective experimental conditions. Intratumoral CD8^+^ T cells after adoptive transfer were sorted as single, live CD8^+^ using a BD FACSAria III cell sorter. Expression of CD45.1/1 (STING-deficient) or CD45.1/2 (STING-expressing) allowed discrimination and sorting according to T cell STING expression. A total of 40,000 cells per condition were loaded per BD Rhapsody cartridge and processed for cDNA synthesis following the BD Rhapsody Single-Cell Capture and cDNA Synthesis protocol (Doc ID: 23-22951(01)).

Library preparation followed the manufacturer’s guidelines outlined in the BD Rhapsody System mRNA Whole Transcriptome Analysis (WTA) and Sample Tag Library Preparation Protocol (Doc ID: 23-24119(02)). Sequencing was conducted on an Illumina NovaSeq X Plus platform following a 150 bp paired-end reads configuration. An average sequencing depth of approximately 50,000 reads per cell was achieved. Data processing, including read alignment, cell barcode demultiplexing, read deduplication, expression matrix generation, and quality reporting, was performed using the Rhapsody Sequence Analysis Pipeline v2.0.

Retrieved count matrices were further analyzed using R v4.3.2. Cells were filtered to include only cells with less than 12.5 % mitochondrial gene content. Multiplets were identified and removed using scDblFinder. The Seurat pipeline v5.1 was used for normalization, dimensionality reduction, and clustering. CD8^+^ T cells were further cleaned using scGate and UCell-based cluster annotation. Gene set enrichment analysis (GSEA) was conducted using Molecular Signatures Database (MSigDB) gene sets.

### Quantification of cGAMP and IFN-**β** by ELISA

For quantification of intra- and extracellular cGAMP in cancer cell lines, cells were seeded in 15-cm culture dishes. For induction of cGAS expression, 1 µg/mL DOX was added. When the cells were 80-90% confluent, the medium was replaced by 5 mL phenol red-free DMEM without additions, except for 1 µg/mL DOX where appropriate. Twenty-four hours later, the conditioned medium was removed and centrifuged at 650 *g* at 4°C for 15 minutes. The supernatant was frozen at –80°C until the assay. The adherent cells were detached with trypsin/EDTA, collected, and centrifuged. Cells were washed once with PBS, centrifuged, and cell pellets were snap-frozen in 1.5-mL reaction tubes using liquid nitrogen for storage at −80°C. Immediately before cGAMP measurement, cell pellets were thawed on ice and lysed using 500 µL M-PER buffer (Thermo Fisher Scientific) per 100 mg cells for 10 minutes, while occasionally vortexing. Cell debris was removed by centrifugation at 14,000 *g* for 15 minutes. For the preparation of tumor tissue, tumor pieces of 100 mg were collected in 500 µL RIPA buffer (Thermo Fisher Scientific) containing proteinase inhibitor (cOmplete, Mini Protease Inhibitor Cocktail, Roche), followed by homogenization with a handheld homogenizer. Homogenized samples were centrifuged at 4°C, 10,000 *g* for 15 minutes followed by collection of the supernatant. Samples were frozen at –80°C until further use. The protein content of lysates was quantified using a DC Protein Assay (Bio-Rad).

The concentration of cGAMP in conditioned media, cell- and tumor-lysates was subsequently measured using a cGAMP ELISA (Cayman Chemical) according to the manufacturer’s instructions.

The concentration of IFN-β was measured using the VeriKine Mouse IFN-Beta ELISA Kit (PBL Assay Science) according to the manufacturer’s instructions. The signal for protein quantification, cGAMP- and IFN-β-ELISA was read on a SectraMAX i3 (Molecular Devices).

### Statistical analysis

One-way or two-way ANOVA was used to compare three or more groups. For comparisons between two groups, either Student’s *t*-test or Mann-Whitney *U*-test was used, depending on data distribution. A *p*-value < 0.05 was considered significant. Statistical analyses were performed using GraphPad Prism software (v9.5.0) or R (v4.3.2).

## Key Resources Table

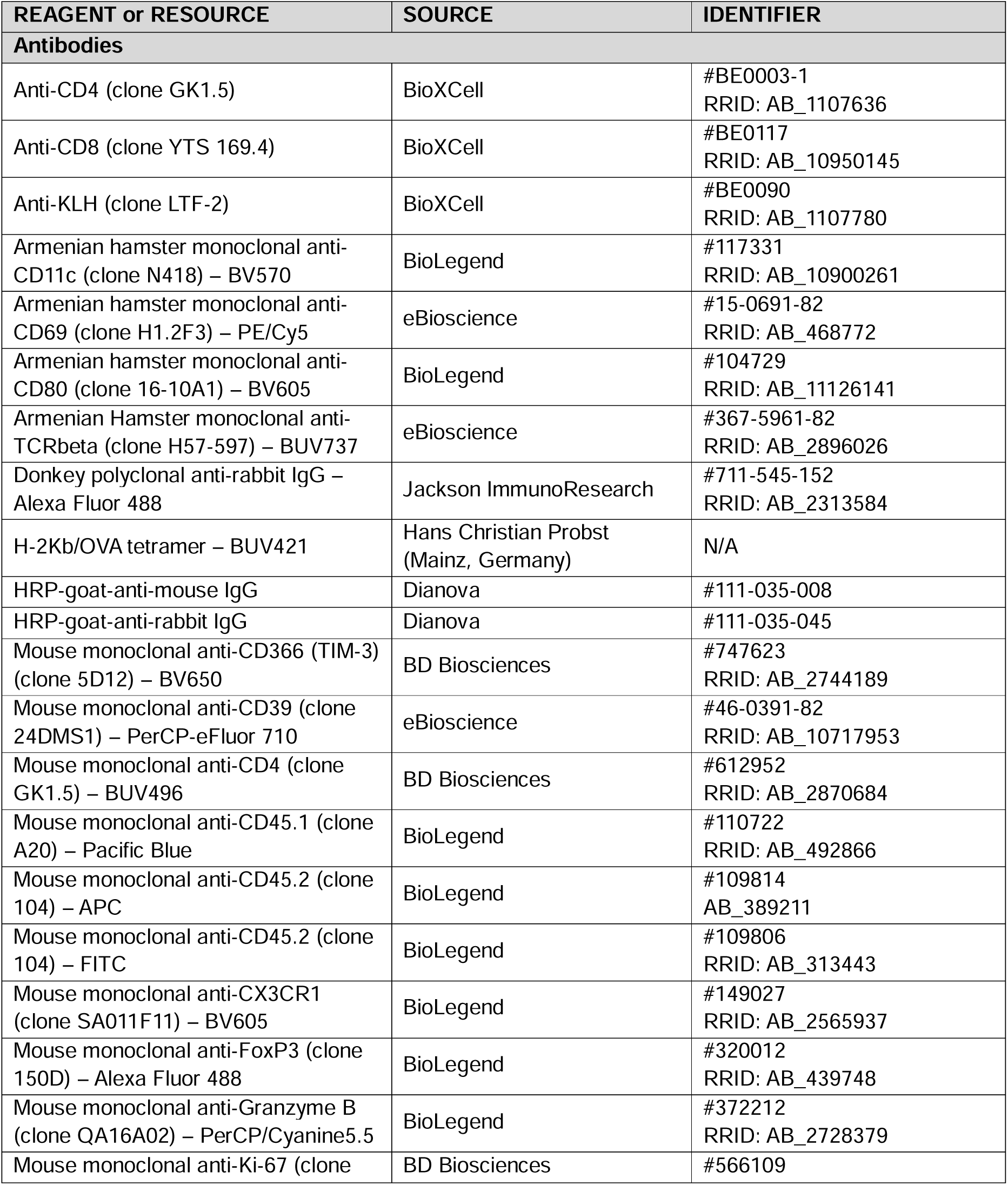
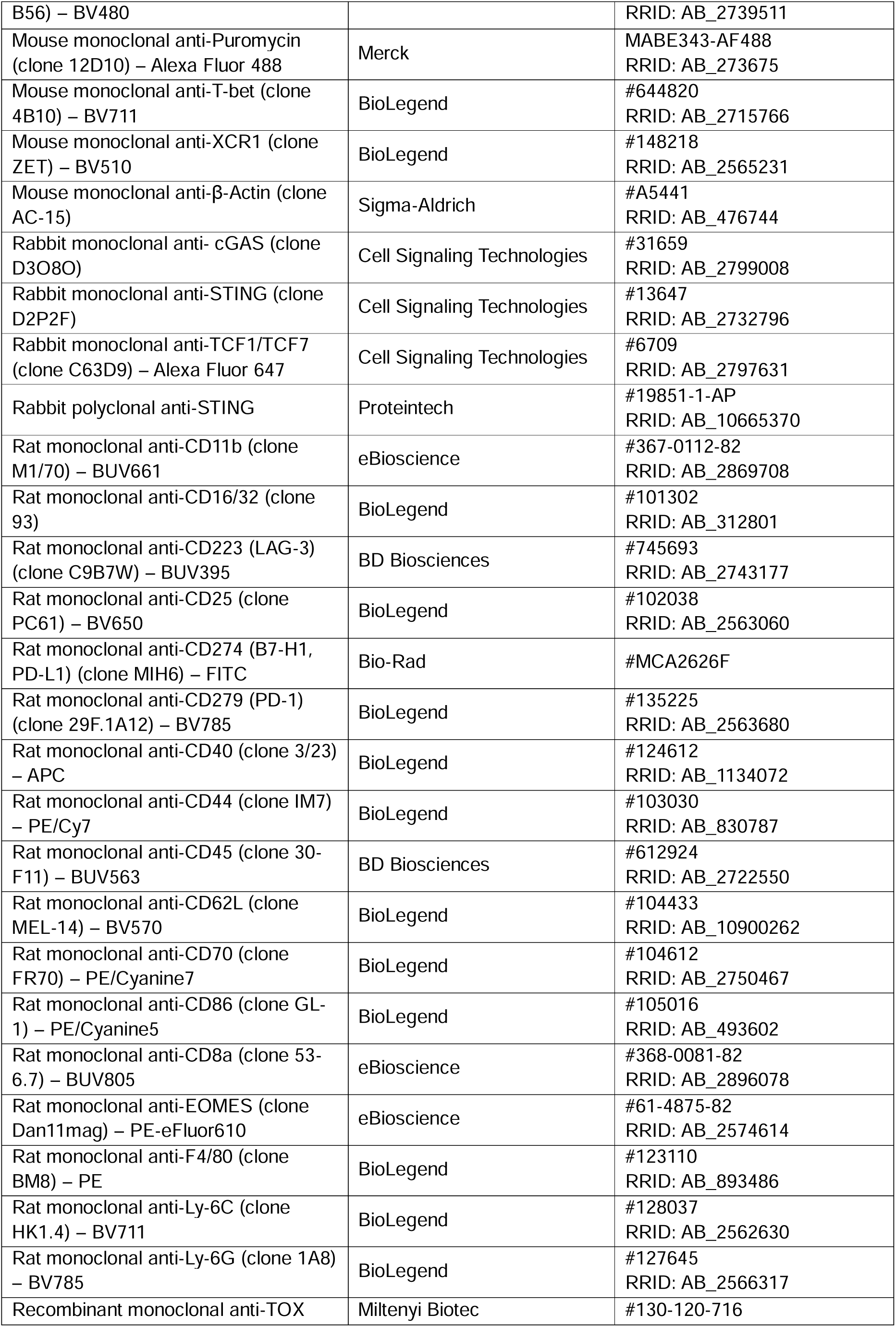
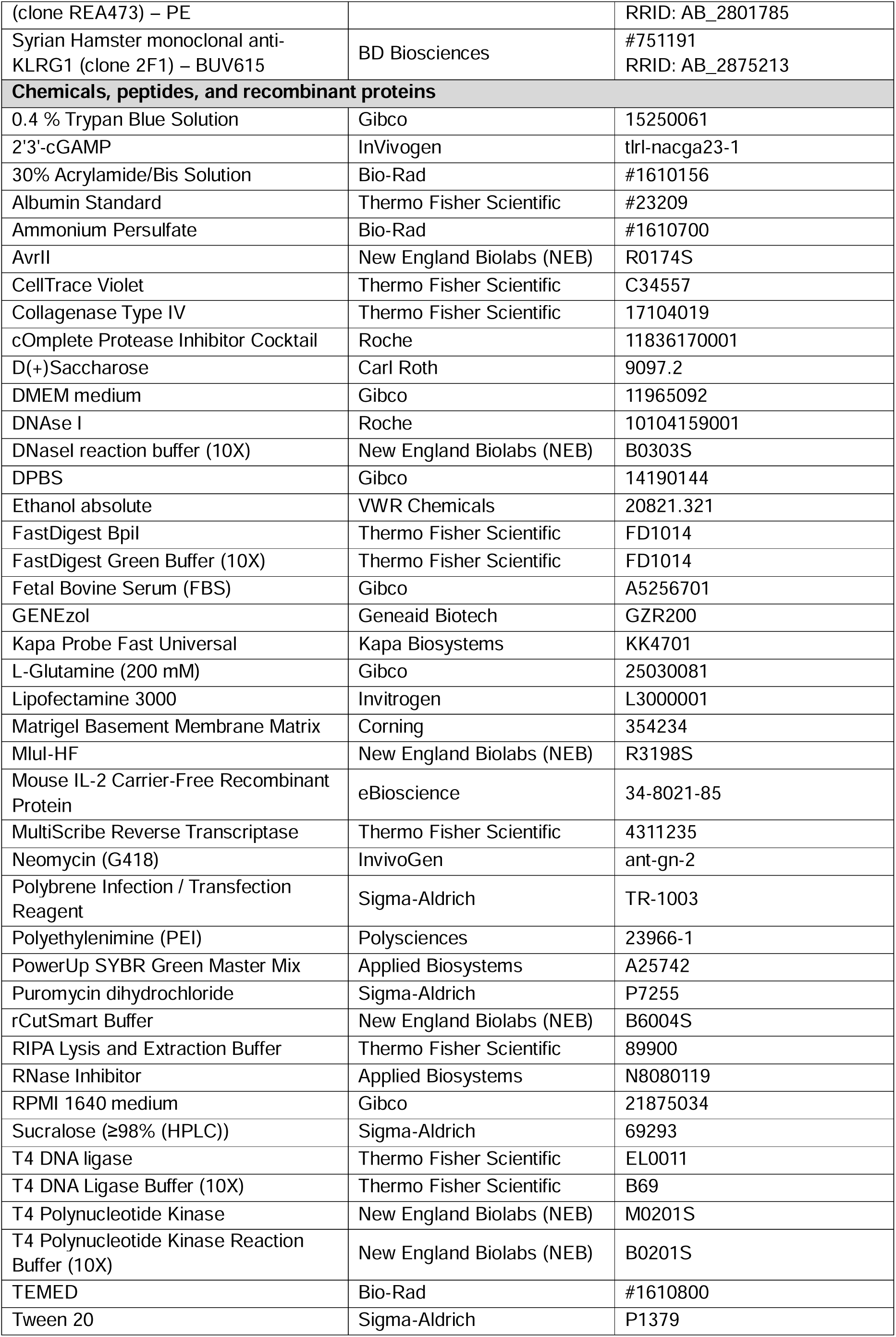
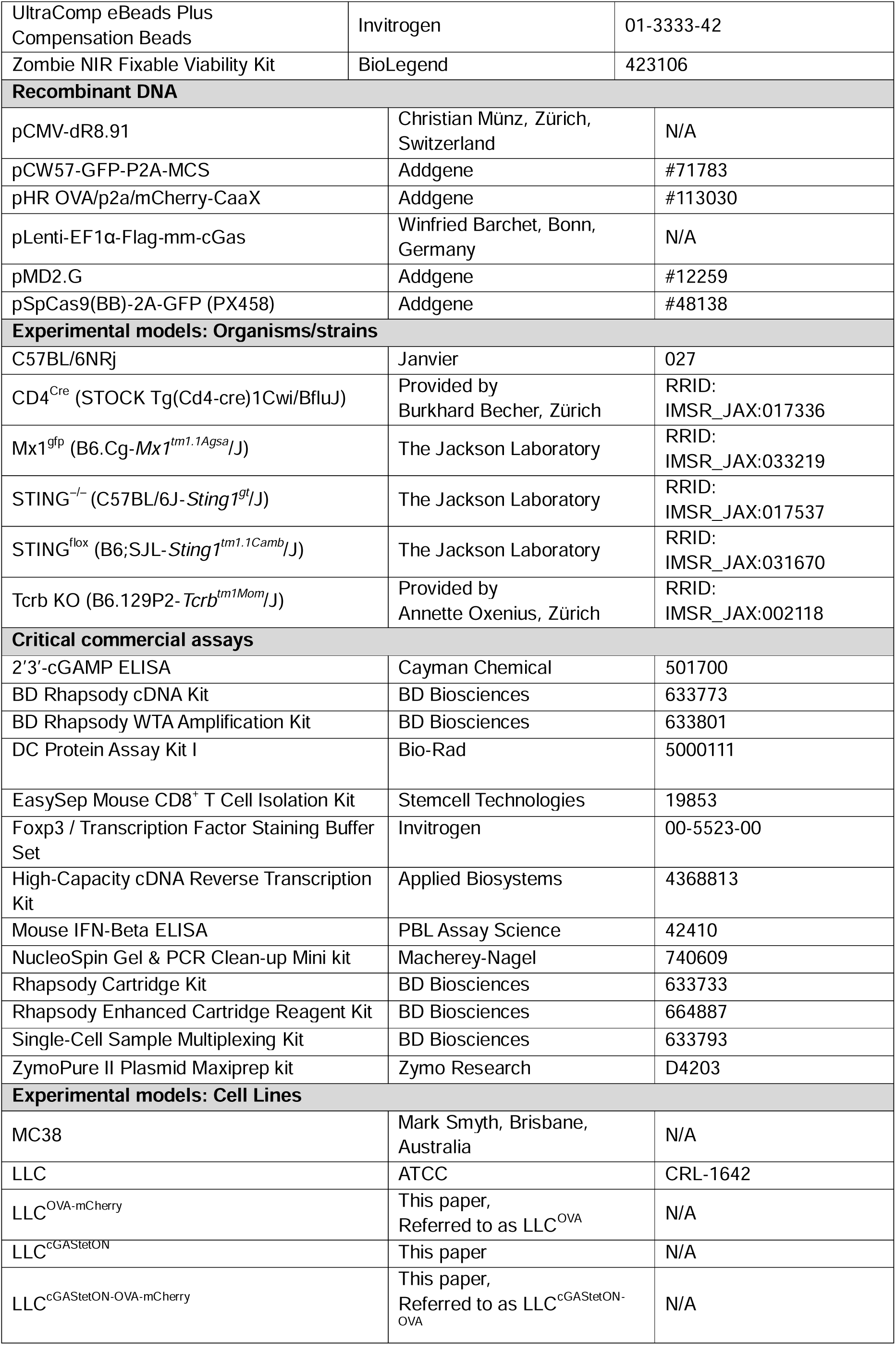
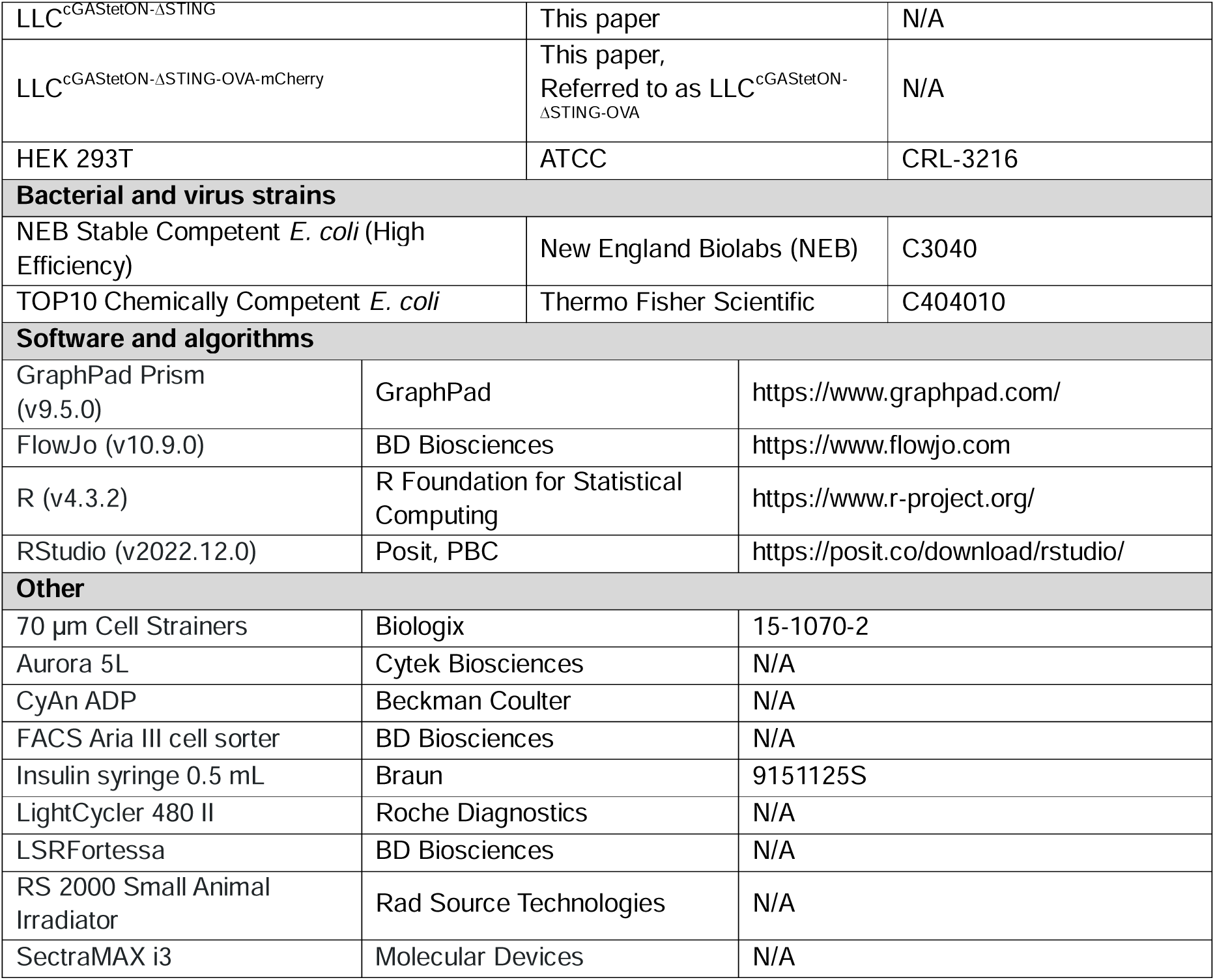

## SUPPLEMENTAL MATERIAL

### Supplemental Methods

#### Quantitative reverse transcription polymerase chain reaction

For gene-expression analysis, 10^5^ LLC^cGAStetON^ cells were seeded in 6-well culture plates and cGAS expression was induced by the addition of 1 µg/mL DOX to the culture medium. Cells were harvested after 4, 8, 16 and 24 hours. For RNA extraction, GENEzol (Geneaid Biotech) reagent was added to the cell layer (500 µL per well) and plates were incubated for 1 minute. Cell lysates were frozen at –80°C until further processing. RNA was isolated by phenol-chloroform extraction. The RNA concentration was measured using a NanoDrop One spectrophotometer (Thermo Fisher Scientific). To remove residual DNA, 10 µg of RNA was treated with DNase I in a 50 µL reaction containing 1X DNase I reaction buffer (NEB) at 37°C for 10 minutes. The reaction was stopped by the addition of EDTA to a final concentration of 5 mM and heat inactivation at 75°C. Reverse transcription (RT) was performed with MultiScribe RT (Thermo Fisher Scientific). SYBR green qRT-PCR was run on a LightCycler 480 II system (Roche). Gene expression was calculated with ΔCt relative to the housekeeping gene *Gapdh*. Primers are listed in **Table S1**.

## Legends to the supplemental figures

**Figure S1.**
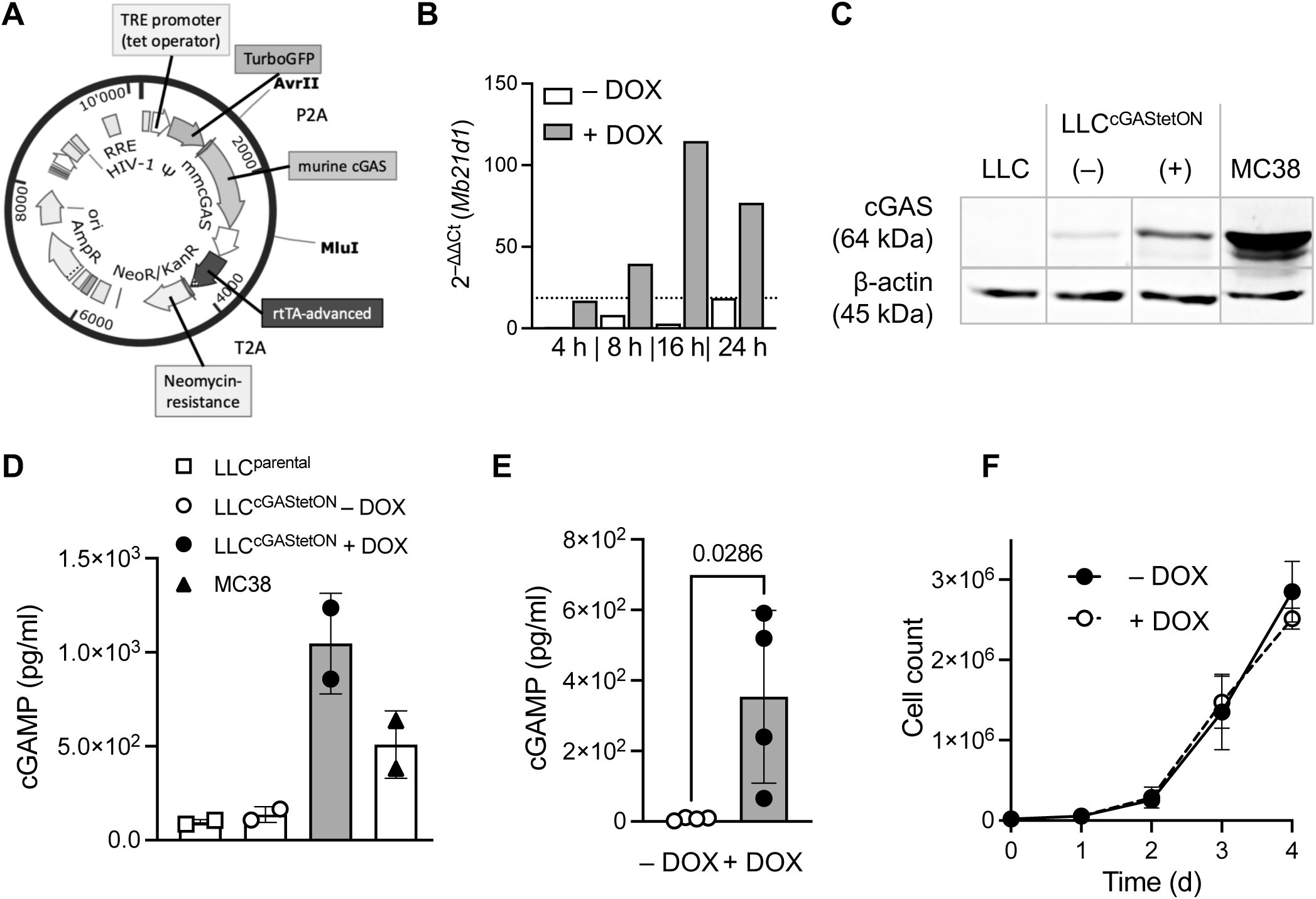
Generation and validation of LLC^cGAStetON^ cells. **A** pCW57-tGFP-P2A-cGAS vector used for the generation of LLC^cGAStetON^ cells. **B** *Mb21d1* transcripts measured in LLC^cGAStetON^ cells 4, 8, 16 and 24 h after induction with 1 µg/mL DOX *in vitro*. ΔCt values calculated relative to *Gapdh* expression. Fold-change calculated as mean ΔΔCt normalized to the 4 h time point without addition of DOX. **C** Western blot showing cGAS expression in LLC, LLC^cGAStetON^, and MC38 cancer cells. LLC^cGAStetON^ cells were cultured with (+) or without (–) 1 µg/ml DOX for 48 h. **D** Quantification of 2′3′-cGAMP in lysates of LLC, LLC^cGAStetON^, and MC38 cells. LLC^cGAStetON^ cells were cultured with or without 1 µg/ml DOX for 24 h. The plot shows two technical replicates per sample. **E** Quantification of cGAMP in cell culture supernatants from LLC^cGAStetON^ cells cultured with or without 1 µg/ml DOX for 24 h. Pooled data from 2 independent experiments are shown. Each symbol represents an individual sample. Statistical analysis: Unpaired two-tailed Student’s t-test. **F** LLC^cGAStetON^ cells were cultured with or without addition of 1 µg/ml DOX and live cells were counted at the indicated time points. Pooled data from 3 independent experiments and 3-4 replicates per time point are shown. The data are expressed as mean ± SD.

**Figure S2.**
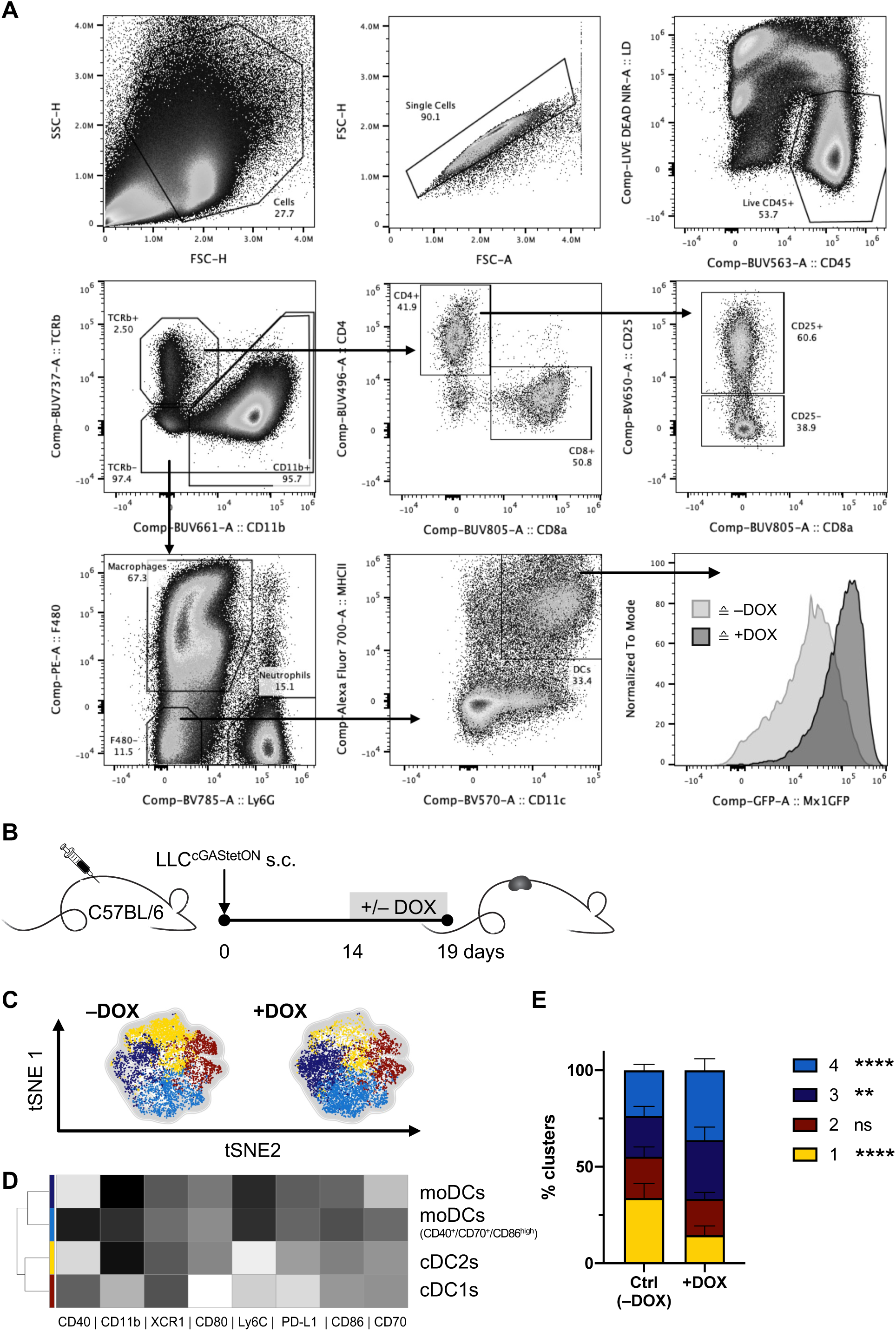
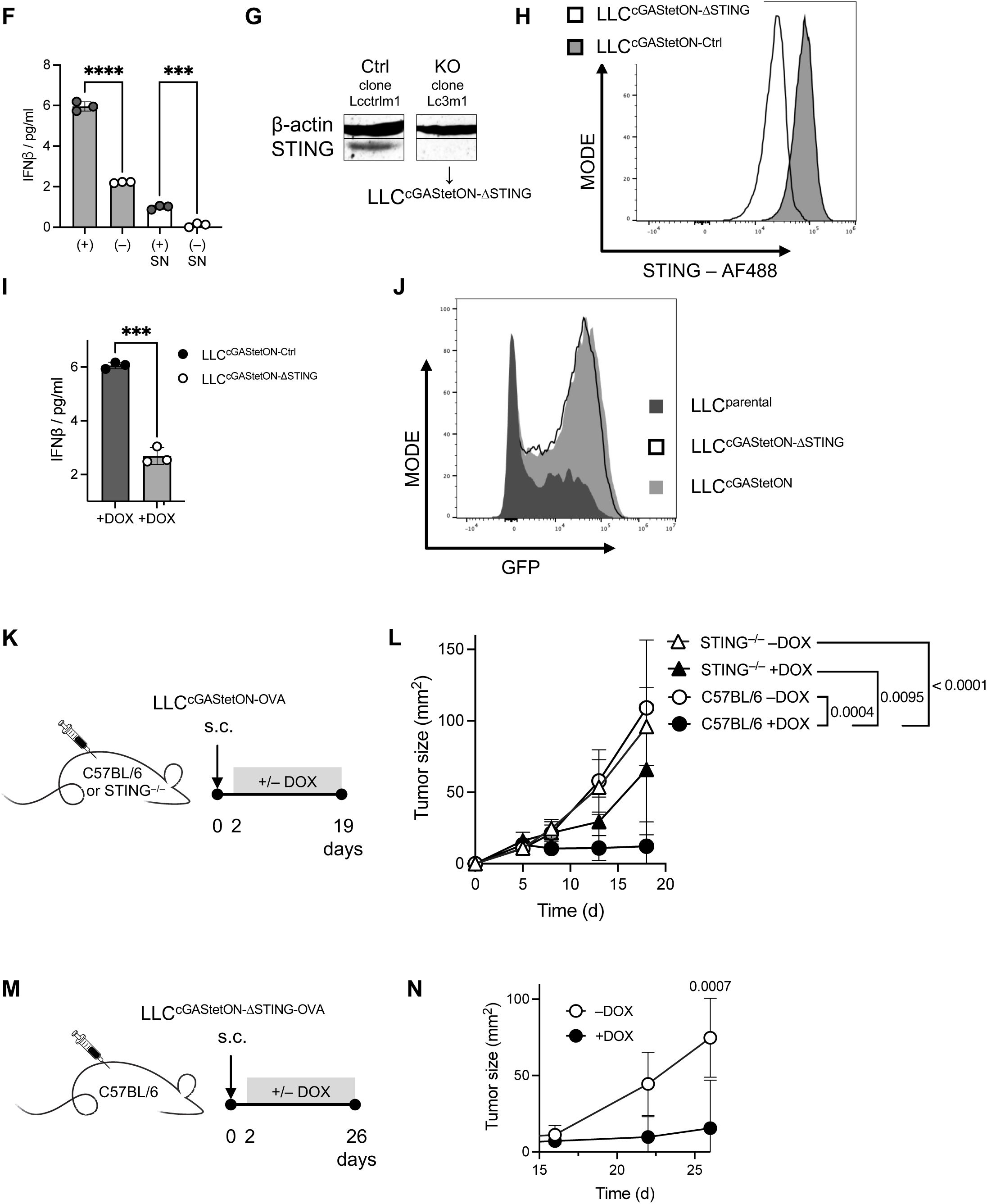
Cancer-cell-intrinsic cGAS modifies the tumor microenvironment and leads to host-STING dependent tumor control. **A** Gating strategy for flow cytometric analysis of intratumoral immune cell populations. **B** Experimental setup for C-E: LLC^cGAStetON-OVA^ cells were s.c. injected into C57BL/6 mice. cGAS expression was induced by DOX from day 14 onwards until the endpoint on day 19. Control mice received no DOX. Results are shown for single, live CD45^+^ TCRβ^-^ F4/80^+^ Ly6G^-^ MHC-II^high^, CD11c^+^ cells (dendritic cells, DCs). **C** tSNE representation of 4 identified DC clusters. **D** Heatmap summary of average expression of indicated markers. **E** Quantification of clusters. Data are plotted as mean ± SD. Statistical analysis: Two-way ANOVA (n = 10 mice per group). **F** IFN-β concentration in cell lysates and supernatant (SN) of DOX (1 μg/ml) treated (+) and untreated (–) LLC^cGAStetON^ cells. The same number of cells (2 x 10^6^) were assayed. The plot shows 3 technical replicates per group. Statistical analysis: Unpaired Student’s t-test. **G** Quantification of STING expression in LLC^cGAStetON-Ctrl^ (Ctrl) and LLC^cGAStetON-ΔSTING^ (KO) cells by Western blot. **H** Representative example showing STING expression in LLC^cGAStetON-Ctrl^ and LLC^cGAStetON-ΔSTING^ cells measured by flow cytometry. **I** Quantification of IFN-β in LLC^cGAStetON-Ctrl^ and LLC^cGAStetON-ΔSTING^ cells by ELISA after 32 h culture with 1 μg/ml cGAMP. Statistical analysis: Unpaired two-tailed Student’s t-test. **J** Representative example showing IFN-I response in LLC, LLC^cGAStetON^ and LLC^cGAStetON-ΔSTING^ tumors measured by flow cytometry. Cancer cells (3 x 10^5^) were s.c. injected in Mx1^gfp^ reporter mice. cGAS expression was induced by DOX from day 10 onwards until the endpoint on day 15. Representative GFP signal in CD4^+^ T cells (single, live CD45^+^ TCRb^+^ CD4^+^ cells). **K** LLC^cGAStetON-OVA^ cells (3 x 10^5^) were s.c. injected into C57BL/6 (n = 10) or STING^-/-^ (n = 9-12) mice. cGAS expression was induced by DOX from day 2 onwards until the endpoint on day 19. Control mice received no DOX. **L** Tumor growth curves. The data are shown as mean ± SD. Statistical analysis: Two-way ANOVA. **M** Experimental setup. LLC^cGAStetON-OVA-ΔSTING^ cancer cells (3 x 10^5^) were s.c. injected into C57BL/6 mice (n = 9 mice per group). cGAS expression was induced by DOX from day 2 onwards until the endpoint on day 26. Control mice received no DOX. **N** Tumor growth. Statistical analysis at the endpoint: Unpaired two-tailed Student’s t-test (n = 10-12 mice per group). The data are expressed as mean ± SD. **p < 0.01, ***p < 0.001, ****p < 0.0001.

**Figure S3.**
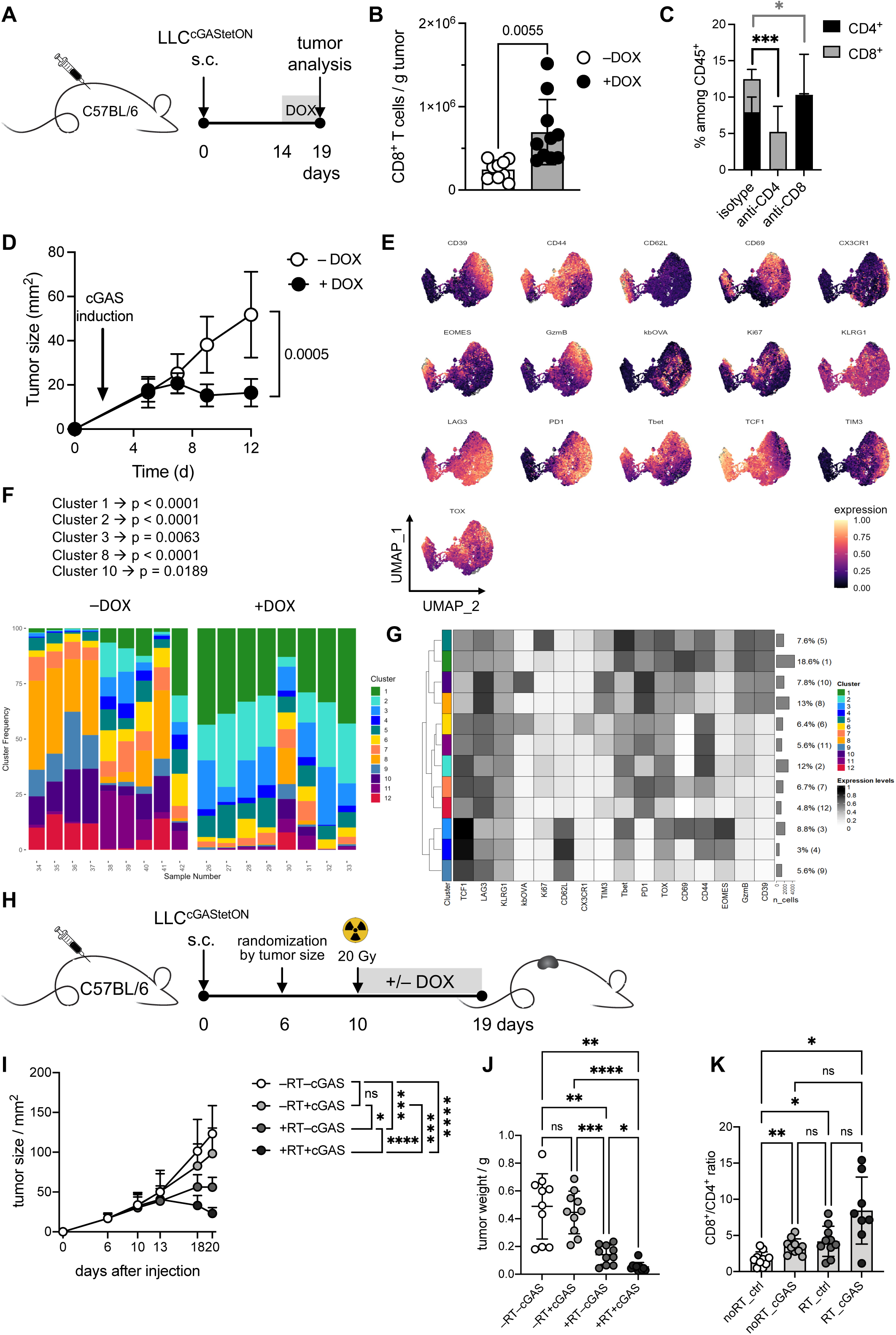
Induced cancer-cell-intrinsic cGAS expression changes frequency and phenotype of tumor-associated CD8^+^ T cells and synergizes with radiotherapy. **A** Experimental setup for B. LLC or LLC^cGAStetON^ cancer cells (2 x 10^5^) were s.c. injected into C57BL/6 mice. Cancer-cell-intrinsic cGAS expression was induced by DOX 14 d later. Control mice received no DOX. **B** Quantification of CD8^+^ T cells in tumors at the experimental endpoint (day 19). Each symbol represents an individual mouse. Statistical analysis: Unpaired two-tailed Student’s t-test. The data are expressed as mean ± SD. **C** Quantification of T cell frequencies in the blood of isotype treated or CD4- or CD8-depleted mice, 5 days after injection of antibodies. Statistical analysis: Unpaired two-tailed Student’s t-test (n = 5 mice per group). The data are expressed as mean ± SD. **D** LLC^cGAStetON-OVA^ cells were s.c. injected into C57BL/6 mice. cGAS expression was induced by DOX from day 2 onwards until the endpoint on day 12. Control mice received no DOX. Plot shows tumor growth as mean ± SD. Statistical analysis at the endpoint: Unpaired two-tailed t-test (n = 9-10 mice per group). The data are expressed as mean ± SD. **E** UMAP dimensionality reduction showing relative expression of functional T cell markers. Analyzed cells gated on single, live CD45^+^ TCRβ^+^ CD8^+^. **F** Graphical display of proportions of identified CD8^+^ T cell clusters in both conditions (–DOX, +DOX). Sample numbers are representative for individual experimental mice. Clusters with statistically significant differences between conditions and respective p-values indicated above. Statistical analysis by unpaired two-tailed t-test with Welch’s correction. **G** Heatmap of relative marker expression per identified T cell cluster. **H** Experimental setup for I-K. LLC^cGAStetON^ (3 x 10^5^) cancer cells were subcutaneously injected in C57BL/6 mice. On day 6, mice were randomized by tumor size and assigned to the different groups. On day 10, tumors were irradiated with 20 Gy. RT, radiotherapy. **I** Tumor growth curves. Statistical analysis: Unpaired two-tailed t-test (n = 10 mice per group). The data are expressed as mean ± SD. **J** Tumor weights at the endpoint (day 19). Each symbol represents an individual mouse. Statistical analysis: One-way ANOVA for data from day 19. The data are expressed as mean ± SD. **K** Ratios of intratumoral CD8^+^ and CD4^+^ T cells, quantified by flow cytometry at the endpoint. Statistical analysis: One-way ANOVA for data from day 19. The data are expressed as mean ± SD. *p < 0.05, **p < 0.01, ***p < 0.001, ****p < 0.0001.

**Figure S4.**
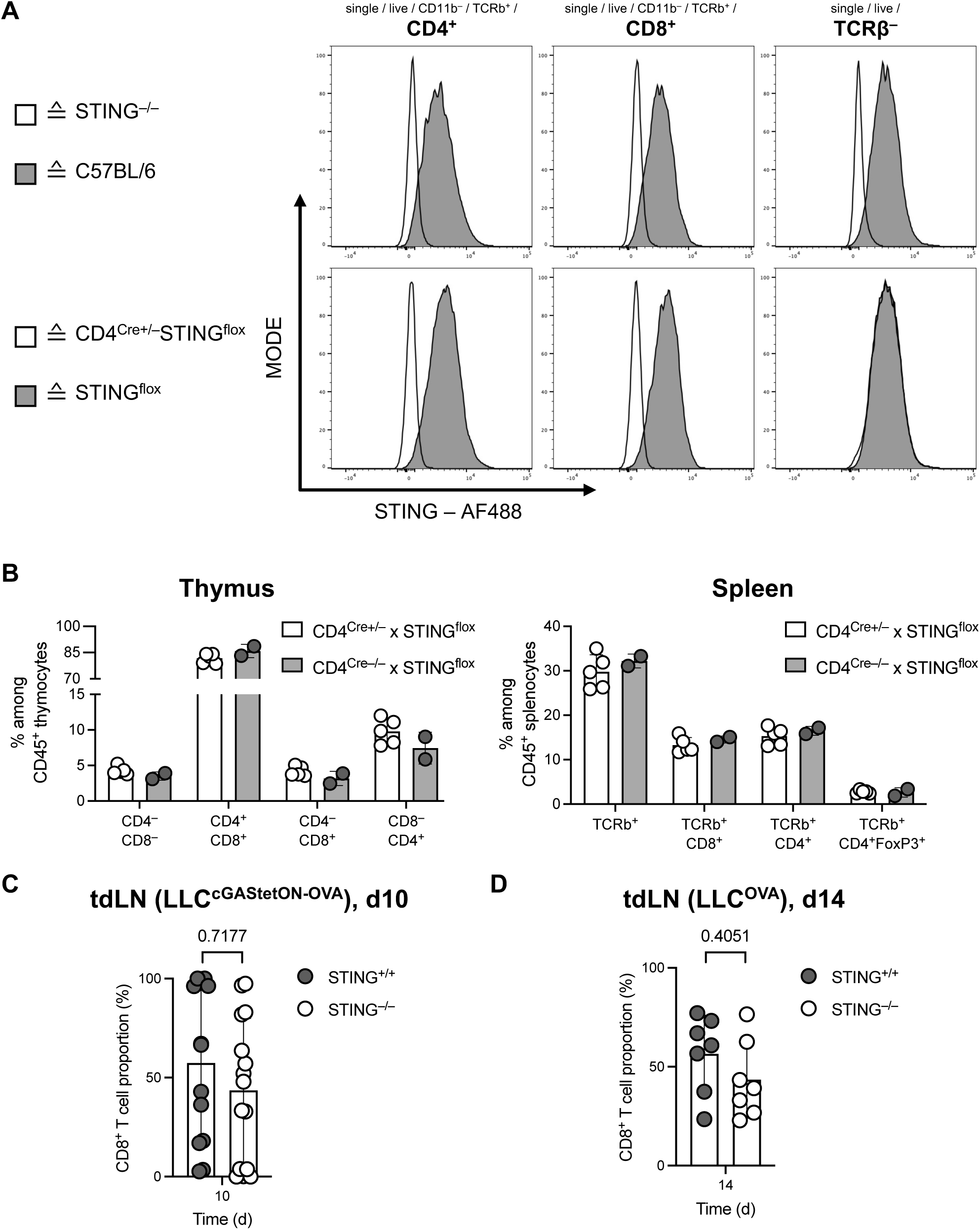
CD4^Cre^STING^flox^ lack STING expression in T cells. **A** STING expression in CD4^+^ and CD8^+^ T cells and TCRβ^-^ cells from naïve C57BL/6, STING-deficient (STING^-/-^), C57BL/6, CD4^Cre+/–^STING^flox^ and STING^flox^ mice was measured by flow cytometry. Representative examples are shown. **B** Proportion of different T cell populations in the thymus (left panel) and spleen (right panel) of CD4^Cre+/–^STING^flox^ and Cre^-/-^ littermate control mice measured by flow cytometry. Statistical analysis: Unpaired two-tailed Student’s t-test. **C** LLC^OVA^ or LLC^cGAStetON-OVA^ cells (3 x 10^5^) were s.c. injected into TCRβ^-/-^ mice. Mice received DOX from day 2 onwards until the endpoint. 3 days after cancer cell injection, 1 x 10^3^ STING^-/-^ plus 1 x 10^3^ STING^+/+^ OT-I CD8^+^ T cells were co-injected via the tail vein. On days 10 and 14, the proportion of transferred T cells were quantified in tdLNs by flow cytometry. Plot shows the proportion of STING^-/-^ and STING^+/+^ OT-I T cells in lymph nodes draining LLC^cGAStetON-OVA^ tumors 7 days after transfer. Each symbol represents an individual sample. Statistical analysis: One-way ANOVA. The data are expressed as mean ± SD. **D** Proportion of STING^-/-^ and STING^+/+^ OT-I T cells in lymph nodes draining LLC^OVA^ tumors 11 days after transfer. Cells were gated on single, live CD45^+^ CD11b^-^ TCRβ^+^ CD8^+^. Each symbol represents an individual sample. Statistical analysis: One-way ANOVA. The data are expressed as mean ± SD.

**Figure S5.**
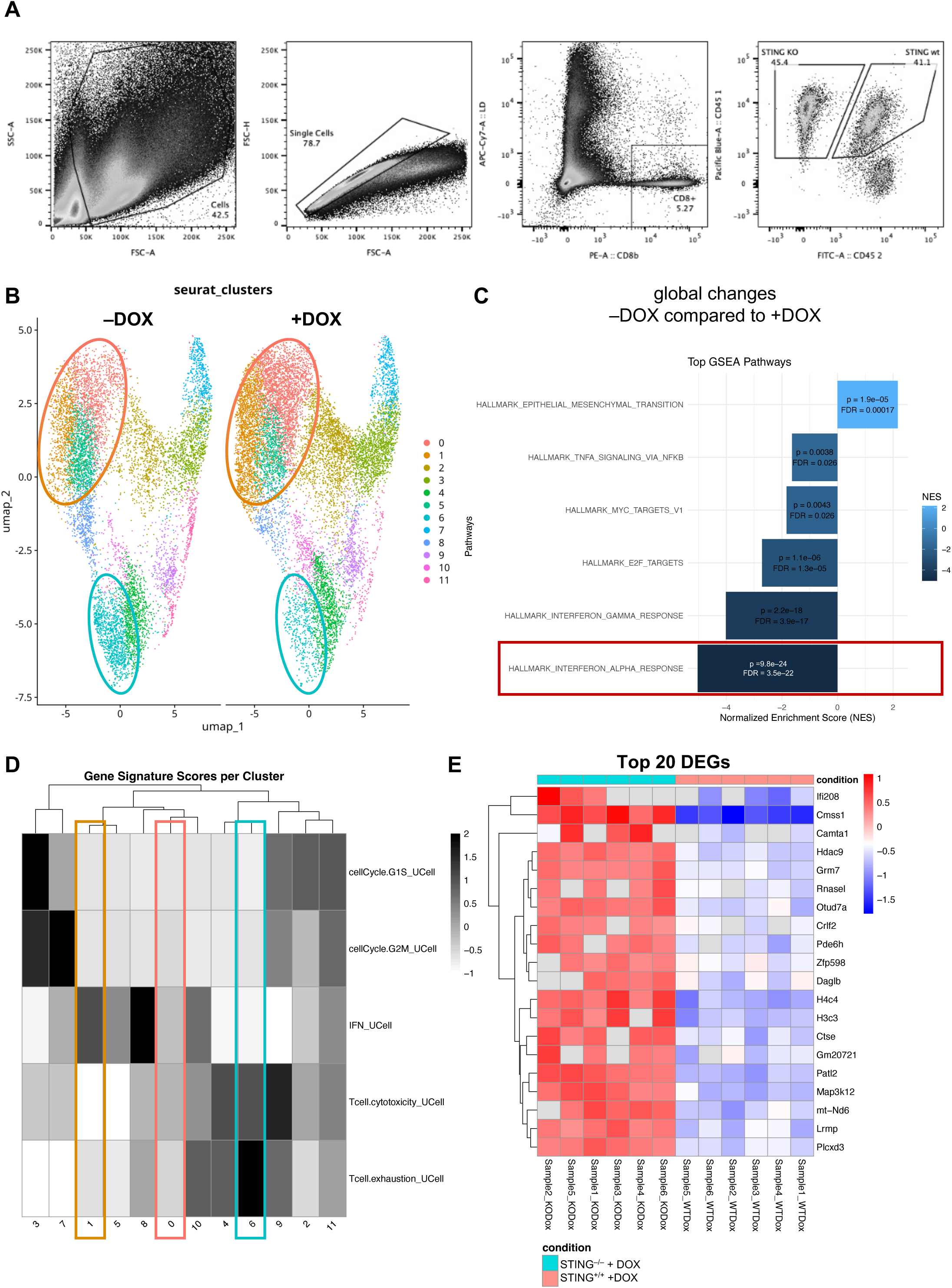
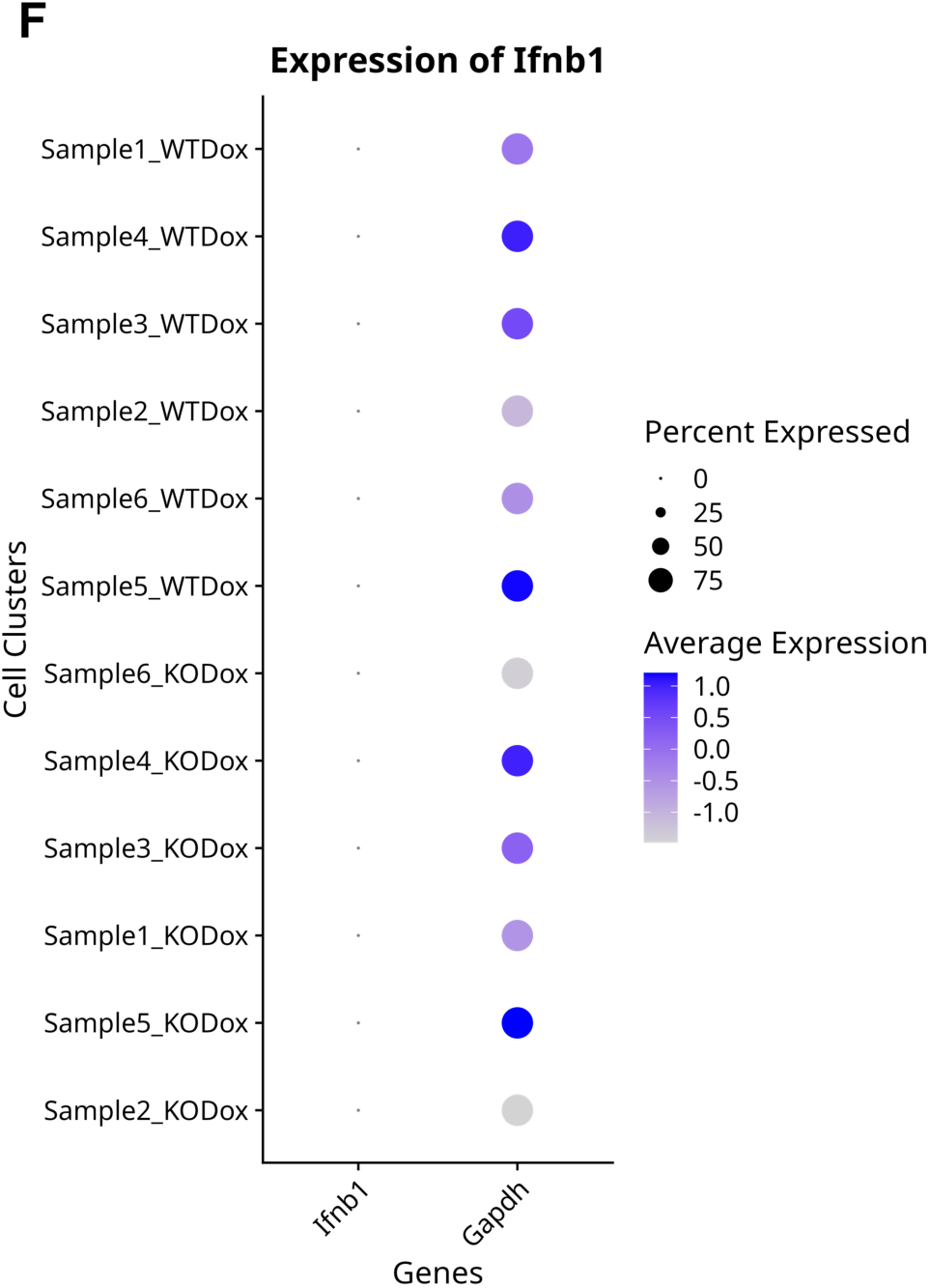
CD8^+^ T cells respond to IFN-I and are affected by cell-intrinsic STING signaling. **A** Sorting strategy for reisolation of CD45.1/45.1 STING^-/-^ and CD45.1/45.2 STING^+/+^ CD8^+^ T cells from LLC^cGAStetON-OVA^ tumors, 6 days after adoptive co-transfer. **B** UMAP representation of 12 (0-11) Seurat clusters comparing STING^+/+^ CD8^+^ T cells with or without induced intratumoral cGAS expression. **C** Gene set enrichment analysis (GSEA) using Molecular Signatures Database (MSigDB) gene sets. **D** Heatmap showing UCell scores for the identified gene signatures across Seurat clusters. **E** Top 20 differentially expressed genes between STING^-/-^ and STING^+/+^ CD8^+^ T cells. **F** Expression of *IFNb1* compared to *Gapdh* per sample.

---

**Table S1:**
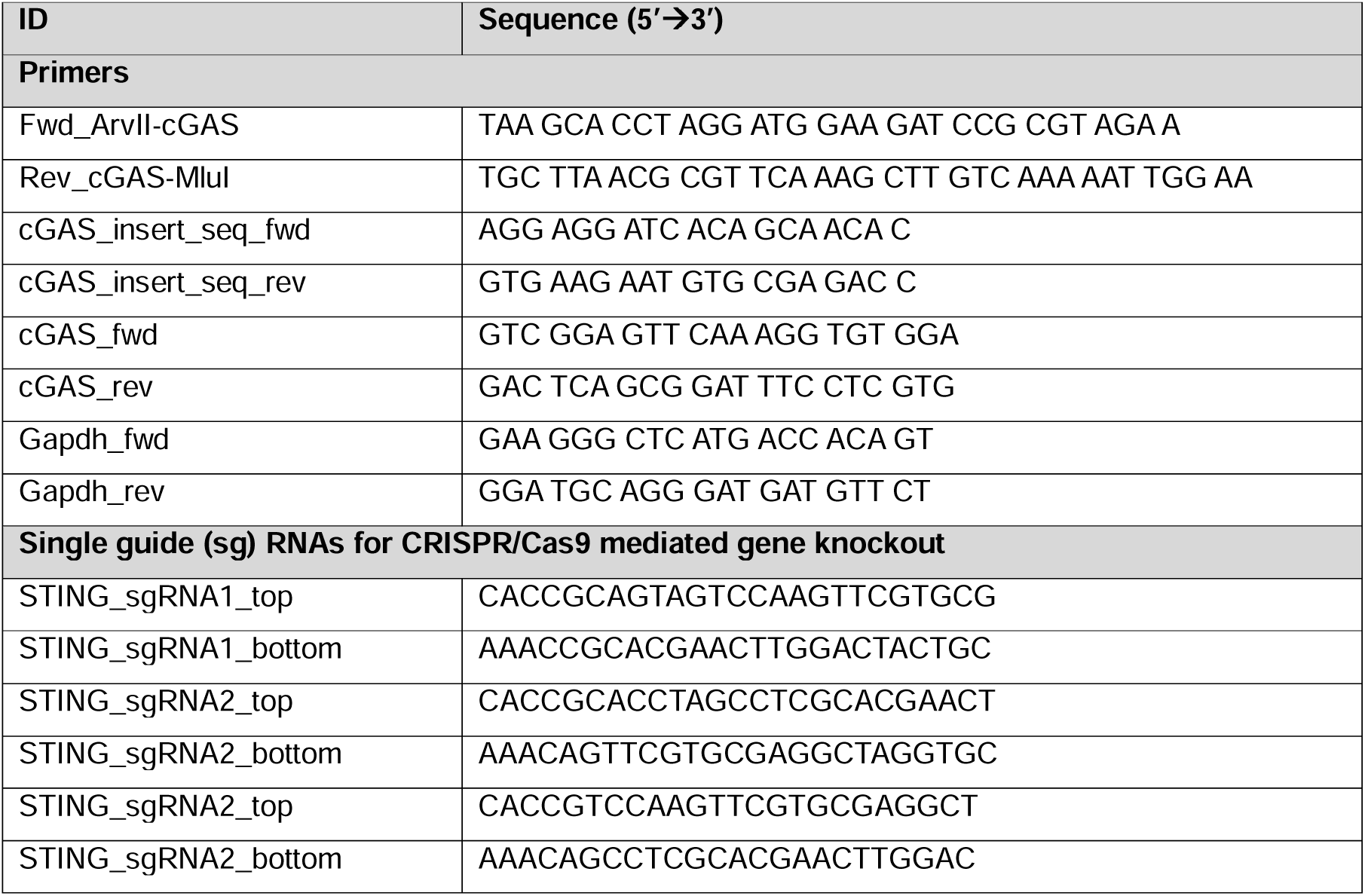
Oligonucleotides.

